# Imbalance and Composition Correction Ensemble Learning Framework (ICCELF): A novel framework for automated scRNA-seq cell type annotation

**DOI:** 10.1101/2024.04.21.590442

**Authors:** Saishi Cui, Sina Nassiri, Issa Zakeri

## Abstract

Single-cell RNA sequencing (scRNA-seq) has gained broad utility and success in revealing novel biological insight in preclinical and clinical investigations. Cell type annotation remains a key analysis task with great influence on downstream interpretation of scRNA-seq data. Traditional machine learning approaches proposed for automated cell type annotation often overlook the inherent imbalance of cell type proportions within biological samples, and the compositional nature of sequencing-based gene expression quantification. In this study, we highlight the importance of accounting for cell type imbalance and compositionality of sequencing count data, and introduce the Imbalance and Composition Corrected Ensemble Learning Framework (ICCELF) as a novel approach to automated cell type annotation. We show via comprehensive evaluation on both simulated and real-world scRNA-seq data that by effectively addressing class imbalance and data compositionality. ICCELF offers a robust and efficient solution that facilitates accurate and reliable cell type annotation, paving the way for enhanced biological discoveries.

## 1. Introduction

Investigating the intricate landscape of gene expression at the single-cell level through RNA sequencing unveils unprecedented insights into cellular heterogeneity and regulatory dynamics, offering a comprehensive understanding of cellular behavior in health and disease. Cell type annotation plays a pivotal role in the analysis of scRNA-seq data as it enables the identification and characterization of distinct cell populations within heterogeneous samples. By accurately assigning cell types to individual cells based on their gene expression profiles, researchers can unravel the cellular composition of tissues and organs, elucidate cell lineage and developmental trajectories, and provide valuable insights into the functional diversity and interactions among different cell types. Moreover, precise cell type annotation is essential for interpreting disease states, identifying novel biomarkers, and developing targeted therapeutic interventions based on scRNA-seq data. However, manual annotation of scRNA-seq data is limited by subjectivity, labor-intensiveness, difficulty in handling cellular heterogeneity, scalability issues, and lack of consistency [1]. To address these limitations, automated methods for cell type annotation have been developed, leveraging computational algorithms and machine learning techniques to streamline the process, improve scalability, and reduce bias, thereby enhancing reproducibility and accuracy in single-cell data analysis [2].

Despite the progress, existing machine learning approaches for automated cell type annotation often overlook the imbalanced and compositional aspects of scRNA-seq data. Single-cell studies generally contain samples with diverse cell types at varying frequencies, leading to imbalanced class distributions. Class imbalance presents a hurdle for statistical learning because most algorithms are built on the premise that the data is evenly distributed within classes. Consequently, models tend to exhibit inferior predictive accuracy, particularly for the minority class. This is suboptimal in applications where the minority class is of greater significance, e.g. representing a rare but clinically relevant cell population. Moreover, akin to other abundance data obtained through next-generation sequencing, scRNA-seq count data exhibits a compositional nature [3]. This implies that the gene abundances observed for each cell are constrained by a fixed total sum referred to as the library size. As the library size is a technical parameter, the abundance of individual genes within a cell holds significance only in relation to other genes in that same cell. Consequently, this constraint dictates that an increase in the abundance of any given gene will inevitably lead to a decrease in the observed abundance of all other genes. These characteristics necessitate normalization of scRNA-seq data prior to application of standard machine learning algorithms for automated cell type annotation. Normalization by library size is the simplest yet most common approach broadly adopted in scRNA-seq data analysis. Despite its simplicity, this method lacks robustness in the presence of differentially expressed genes [4]. Although more sophisticated normalization methods have been proposed [5], they are generally not compatible with machine learning applications where cells ought to be analyzed one at a time.

## 2. Materials and Methods

### 2.1 Data

To validate our approach, we utilized two types of datasets in our analyses: real-world scRNA-seq data from peripheral blood mononuclear cells (PBMCs), and simulated scRNA- seq data with controlled parameters. The real-world PBMC datasets were sourced from 10X Genomics. These datasets specifically include 2,700 PBMCs from the CITE-seq reference data set, which was recently published in a study by Hao et al [6].This reference data set consists of 162,000 PBMCs measured using 228 antibodies. The benchmarking dataset includes 24 human PBMC data sets, representing 8 different donors and 3 different batch times. These datasets were manually annotated at three different resolutions: fine, moderate, and coarse annotation. The fine annotation contains 56 cell types, but due to some cell types having zero cells, we decided to focus on the moderate and coarse annotations for our working datasets. The 24 datasets with coarse annotation labels were considered as our real- world coarse-resolution datasets. These datasets consist of 8 different cell types: B cells, CD4 T cells, CD8 T cells, dendritic cells (DCs), monocytes (Mono), natural killer cells (NK cells), Other cells, and Other T cells. For our fine-resolution datasets, we used the moderate- resolution labels but restrict them to T cells. This results in 24 fine-resolution datasets containing 11 types of T cells: CD4 cytotoxic T lymphocytes (CTLs), CD4 naïve T cells, CD4 central memory T cells (TCMs), CD4 effector memory T cells (TEMs), CD8 naïve T cells, CD8 TCMs, CD8 TEMs, double-negative T cells (dnT), gamma-delta T cells (gdT), mucosal-associated invariant T cells (MAIT), and regulatory T cells (Tregs). The summary statistics of these 24 PBMC datasets can be found in Table S1. Without loss of generality, within each resolution we trained on the first dataset and predicted on the other 23, emulating real-world scenarios with only one training set, because obtaining multiple fully annotated sets for integration is challenging, and integration can introduce noise or bias. The specific first set choice for training is arbitrary, as permutation does not matter. As we train the machine learning model on the first dataset for each resolution, we can apply the trained model to the remaining testing sets in the same donor (different batches) to evaluate performance within the same donor. Additionally, applying the model to testing sets across different donors provides results across donors with the same trained model. This allows us to assess performance both within the same donor using different batches as well as across different donors.

For the simulation study, we designed 5 scenarios to generate datasets with varying levels of class imbalance, from highly imbalanced to moderately imbalanced to balanced. This enables assessment across different imbalance conditions while controlling other factors. Table 1 displays the distribution of cell types for each scenario. Both the training and testing sets contain 6,000 cells in total. Scenarios 1-5 represent decreasing imbalance, as evidenced by the decreasing standard deviation of cell type samples in Table 1. Scenario 5 and the testing data are completely balanced. For each scenario, 1 training set was simulated with a predefined imbalance level. Then 20 testing sets were generated using different random seeds for final evaluation per scenario. The simulated datasets were generated using the SPARsim [7] package in R [8], which creates count matrices resembling real data by modeling the distribution of zeros with a Gamma-Multivariate hypergeometric distribution. In evaluations by Crowell et al. [9] and Cao et al. [10], SPARsim emerged as one of the top performing scRNA-seq simulators, closely mimicking real data properties. To set the SPARsim parameters, we estimated them from the 10X Genomics example datasets of human Jurkat and 293T cells from Zheng et al. The 293T sample of 1,718 cells is included as a built-in SPARsim dataset. Using the estimated 293T parameters, we simulated 10 cell groups with abundances defined in Table 1 and fixed 15,000 total genes for each scenario. Within each cell group, 20 “driver” genes were designated that are uniquely upregulated for that cell type, totaling 200 driver genes. We also selected 1,800 “DE” (differentially expressed) genes SPARsim uses a fold change multiplier to control differential expression, with >1 indicating upregulated or downregulated in 2+ cell types. This resulted in 2,000 informative genes. upregulation and < 1 indicating downregulation. For upregulated DE genes, the multipliers were sampled from *unif(a_up,_ b_up_*) where *a_up ∼_ N*(1.8, 0.025) and *b_up ∼_ N*(2.0, 0.025). For downregulated DE genes, the multipliers were sampled from *Unif(a_down,_* b*_down_*) where *a_down ∼_ N*(0.3, 0.025) and *b_down ∼_ N*(0.5, 0.025). For driver genes, the upper bounds were multiplied by 1.2. The designation and parameter settings for highly variable genes in the simulated datasets are provided in Table S2. These designations and parameters are kept consistent across all 5 scenarios in order to generate controlled datasets with predefined class imbalance or balance.

**Table 1.** Distribution of simulated cell types for each scenario.

|  | Training datasets |  |  |  |  | Testing datasets |
| --- | --- | --- | --- | --- | --- | --- |
| N (%) | Scenario 1 | Scenario 2 | Scenario 3 | Scenario 4 | Scenario 5 |  |
| Cell type 1 | 60 (1) | 120 (2) | 240 (4) | 360 (6) | 600 (10) | 600 (10) |
| Cell type 2 | 180 (3) | 240 (4) | 360 (6) | 420 (7) | 600 (10) | 600 (10) |
| Cell type 3 | 240 (4) | 360 (6) | 360 (6) | 420 (7) | 600 (10) | 600 (10) |
| Cell type 4 | 360 (6) | 480 (8) | 480 (8) | 540 (9) | 600 (10) | 600 (10) |
| Cell type 5 | 420 (7) | 480 (8) | 480 (8) | 600 (10) | 600 (10) | 600 (10) |
| Cell type 6 | 480 (8) | 600 (10) | 600 (10) | 600 (10) | 600 (10) | 600 (10) |
| Cell type 7 | 600 (10) | 600 (10) | 600 (10) | 600 (10) | 600 (10) | 600 (10) |
| Cell type 8 | 960 (16) | 840 (14) | 720 (12) | 720 (12) | 600 (10) | 600 (10) |
| Cell type 9 | 1,200 (20) | 960 (16) | 900 (15) | 780 (13) | 600 (10) | 600 (10) |
| Cell type 10 | 1,500 (25) | 1,320 (22) | 1,200 (20) | 840 (14) | 600 (10) | 600 (10) |
| Total | 6,000 (100) | 6,000 (100) | 6,000 (100) | 6,000 (100) | 6,000 (100) | 6,000 (100) |
| SD (Mean, SD) | 471.6 (7.9) | 357.8 (6.0) | 281.4 (4.7) | 154.9 (2.6) | 0 (0) | 0 (0) |

### 2.2 Workflow of ICCELF

The schematic of ICCELF is shown in Figure 1. First, we obtain a scRNA-seq dataset with annotated true labels as training set, which is imbalanced with different cell type proportions. We then employ our proposed imbalance correction technique described in 2.3, oversampling minority classes and downsampling majority classes. For the downsampled data, we directly apply centered log-ratio (CLR) transformation described in 2.4 due to compositional coherence. For the oversampled data, we take the CLR of original minority samples and synthesize new samples by concatenating with CLR of oversampled data. For example, if an original minority class has 60 samples, oversampled to 500, the CLR composition is:

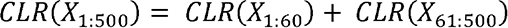

**Figure 1.**
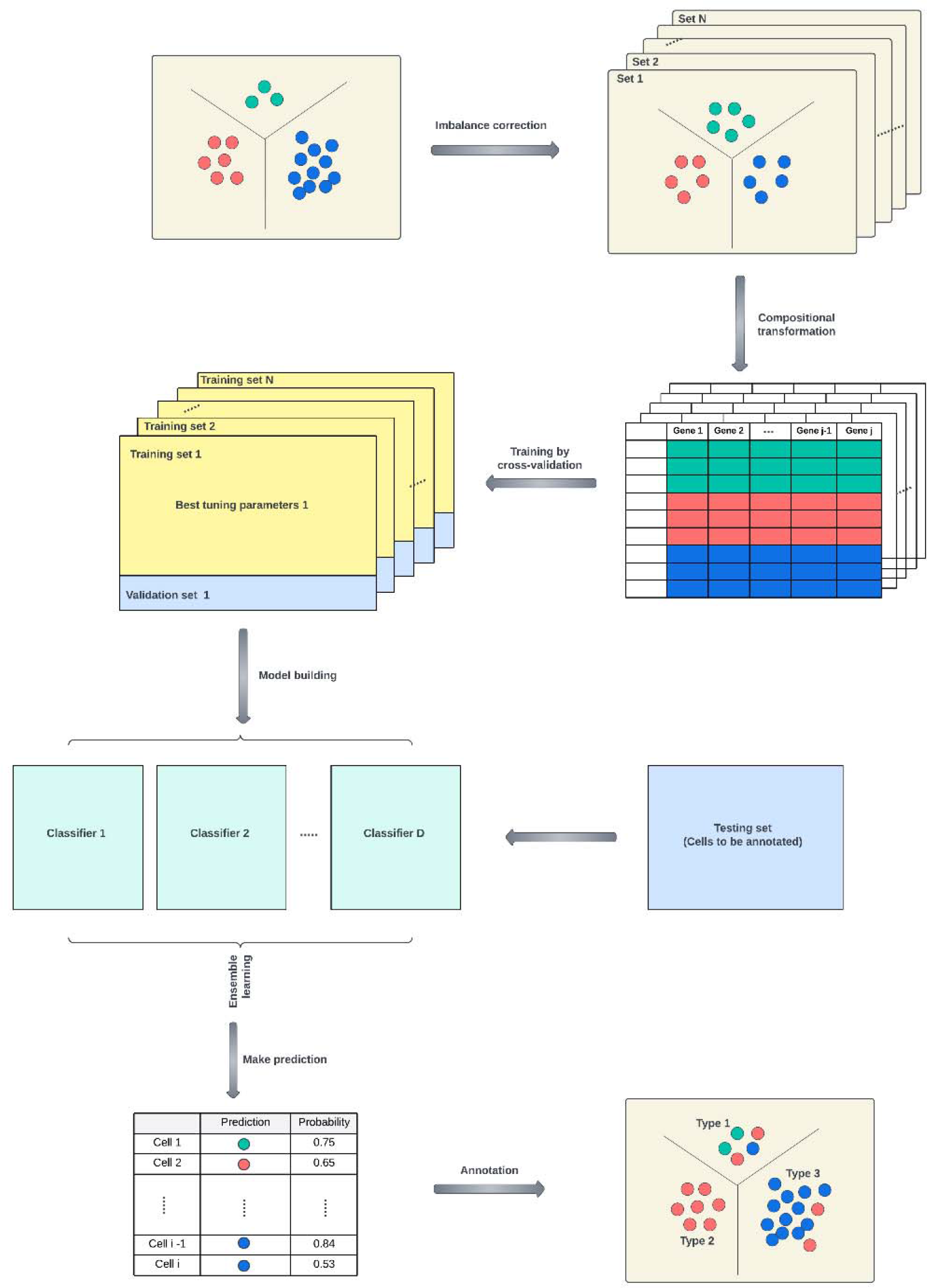
Schematic of ICCELF workflow. The framework consists of four major steps: 1). Imbalance correction 2). Log-ratio (compositional) transformation 3). Machine learning classifiers training 4). Ensemble the predictions and make annotations.

where, *X*_1:60_ is the original sample of a minority cell type, *X*_61:500_ is the oversampled one, and, *X*_1:500_ is the final synthetic sample. We create .5training datasets (layers) with this procedure. For each layer, we train the machine learning classifiers described in 2.5, evaluating both multi-class and binary performance to determine the best approach. For prediction, we take CLR of raw test data, apply classifier-specific transformations, and predict with each of the 5 models to get probabilities 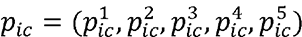 for cell *i*, and class *c*. We ensemble by averaging these 5 probabilities to get final prediction *p_ic_*.

### 2.3 Dealing with class imbalance

Class imbalance is a prevalent challenge in classification tasks, both binary and multiclass. This imbalance occurs when one class is significantly more represented than others, potentially affecting the accuracy of models. This issue is seen in diverse areas such as fraud detection, spam filtering, disease screening, and even in automatic cell annotation in scRNA-seq data analysis. The impact of class imbalance on training and testing machine learning classifiers is a critical area of focus. It involves multiple aspects that need careful examination to mitigate any negative effects. One key aspect is the phenomenon of neutral training, where classifiers become biased towards the majority class. This bias stems from the tendency of many algorithms to prioritize overall accuracy, often neglecting the minority class. Consequently, the underrepresented class may be ignored, leading to significant performance disparities in the classifier’s results. Another consequence of class imbalance is poor generalization. Despite its exceptional performance on the majority class, a classifier may struggle to generalize well for the minority class. Additionally, due to biased training and poor generalization, class imbalance may result in misleading performance metrics. Traditional metrics like accuracy can be deceptive in the presence of imbalanced classes. To illustrate this, let’s consider a scenario where 95% of the information is linked to Category A and only 5% to Category B. In such a situation, even a basic classifier that labels everything as Category A will still achieve a 95% accuracy. However, this accuracy is misleading because the classifier is completely failing to recognize Class B instances. This emphasizes the significance of utilizing alternative performance metrics that are more responsive to the performance of the minority class, such as precision, recall, or F1 score. On the whole, the influence of class imbalance in training and testing machine learning classifiers should not be overlooked. It has the potential to introduce significant biases, hinder generalization, and mislead performance evaluation. As such, it is crucial for researchers to tackle this matter and devise effective strategies to alleviate the adverse impacts of class imbalance. These strategies may include data resampling techniques, algorithmic modifications, or the use of specialized performance metrics. By performing this action, we can guarantee that machine learning classifiers are just, strong, and able to precisely handle unbalanced datasets.

One technique to address class imbalance is random undersampling, where observations from the majority class are selectively removed until class equilibrium is reached. While useful when abundant data exists, undersampling risks losing valuable information. Another prevalent approach is oversampling the minority class. Random oversampling simply replicates minority samples, which doesn’t add new information and can cause overfitting. Synthetic Minority Oversampling Technique (SMOTE) [11] instead generates synthetic minority data. SMOTE focuses on the minority class. For each minority sample, it chooses *k*, Synthetic Minority Oversampling Technique (SMOTE) [11] instead generates synthetic nearest neighbors. It randomly selects one neighbor and calculates the feature vector difference between that sample and neighbor. This difference is multiplied by a random number between 0 and 1 and added to the original sample’s feature vector to synthesize a new sample. While SMOTE and its variants have benefits, these oversampling techniques havemethods, overfitting to training data is possible; 3). the choice of, is subjective, giving limitations: 1). excessive oversampling can introduce noise; 2). like other oversampling inconsistent results; 4).,-NN struggles in high dimensions, especially with highly multi-dimensional scRNA-seq data. In summary, oversampling techniques like SMOTE aim to address class imbalance but have notable drawbacks to consider, including overfitting and poor performance on complex high-dimensional datasets.

Consequently, it is vital to devise an innovative oversampling technique specifically formulated for scRNA-seq data. In this study, we employ the concept of testing Poisson a scRNA-seq dataset are independent, where *Xcj*, represents the gene expression values of cell variance [12], with further elaboration provided in Appendix A. We assume that the genes in type, (which is considered a minority class requiring oversampling) and gene,. It is well-established that the expression values in scRNA-seq follow either a Poisson or negative binomial distribution [13, 14], depending on the presence of overdispersion. By employing the theorem in Appendix A, we can ascertain whether a gene exhibits overdispersion or not. There are three situations: if the expression values for gene, in cell type, exhibit overdispersion, synthetic values are generated using a negative binomial distribution; if there values are detected for gene, in cell type,, zero values are generated. is no evidence of overdispersion, a Poisson distribution is employed; and if no expression values are detected for gene *j* in cell type *C*, zero values are generated.

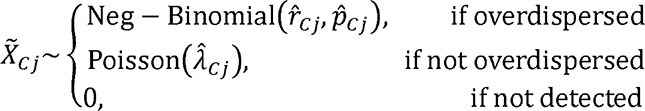

Where *X_cj_* are i.i.d. oversampled expression values for minority class 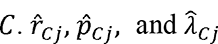 are the method of moments estimates of distribution parameters. Specifically,

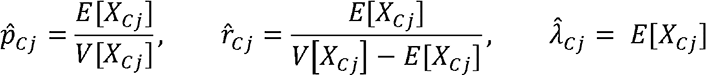

Where *E*[*X_cj_*] and *E*[*X_cj_*] are the mean and the variance of the original sample of minority class *C*. After developing the oversampling technique for scRNA-seq data, we integrate random undersampling in order to create an imbalance correction strategy. Assuming we have a training set of scRNA-seq data, containing *d* annotated cell types, where the sample size are {*S_1,_* …, *S_d_*} the objective of this strategy is to achieve data set balance. We find the median of the sample size, denoted as *S_d._* The cell types with sample sizes less than *S_d_* will undergo oversampling using our proposed approach, while the cell types with sample sizes greater than *S_d_* will undergo random undersampling. As mentioned earlier, random undersampling of the majority class can result in the loss of valuable information. Hence, we will repeat this process of oversampling and undersampling to generate *B* (typically 5-10) training layers. In the subsequent steps, we will train machine learning classifiers on these *B* training data layers and then ensemble the predictions.

### 2.4 Compositional data transformation

As mentioned above, the integration of NGS data analysis can be effectively realized by recognizing the compositional nature of such data. High-throughput sequencing (HTS) count data are inherently compositional, meaning their true significance emerges when considered in terms of relative abundances. Compositional data views each sample as a composition, represented by a vector of non-zero positive values, or components, that convey relative information [15]. This type of data possesses two distinct characteristics. Firstly, the aggregate of all component values, known as the library size, is a byproduct of the sampling method [16]. Secondly, the variance between component values is interpretable only in proportional terms. For instance, a disparity between 100 and 200 counts conveys identical information to a difference between 1,000 and 2,000 counts [17]). Compositional data reside not in the conventional Euclidean space but in a specialized sub-space termed the simplex [15]. However, several frequently employed metrics mistakenly operate under the assumption of them existing in the former space. Such metrics prove unsuitable for relative data, encompassing distance measures, correlation coefficients, and multivariate statistical models [18]. Specifically, the distance between any two variables within compositional data becomes unpredictably affected by the presence or absence of other components [19]. Furthermore, multivariate statistics can produce misleading outcomes, as depicting variables as fractions of a whole inherently makes them interdependent. For example, increasing one variable’s abundance proportionally reduces that of others [20]. This concept equally applies to NGS abundance data [21]. By transforming data into real space using log-ratio transformation, the closed compositional data become open, measurements such as Euclidean distance become meaningful, and a variety of statistical methods can be applied, both for unsupervised and supervised learning, on a regular interval scale [20]. Several log-ratio transformations exist for handling compositional data [21]. Common approaches include: 1). Pairwise and additive log-ratios; 2). Centered log-ratios (CLR); 3). Log-contrasts; 4). Isometric log-ratios (ILR). In this study, we utilize CLR for its conceptual and computational ease. The CLR transformation is a symmetric transformation of the parts *x_j_*(*j* ɛ {1, …, *D*}) (*x_j_* is a vector of expression value of gene *j*), defined as the log-ratios of individual parts with respect to their geometric mean *g*(*x*) = (*x_1,_ x_2,_* …, *x_D_*)^1/*D*^:

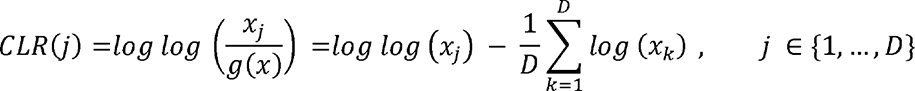

Once the data matrix is transformed using log-ratios, it satisfies four crucial principals of compositional data analysis: 1). Scale Invariance: Multiplying a composition by a constant factor doesn’t alter the results. 2). Perturbation Invariance: Changing units of a composition to equivalent units doesn’t affect the results. 3). Permutation Invariance: Altering the order of components in a composition doesn’t change the results. 4). Sub-compositional Dominance: A subset of the complete composition carries less information than the whole. Moreover, these ratios can be treated as if they were unconstrained, transitioning from closed and constrained to open and unconstrained data. This transformation into real space makes measurements like Euclidean distance meaningful, enabling the application of multivariate statistical methods, as well as machine learning and deep learning techniques. In later of the paper, our study will delve into various machine learning methods for classification, with these features are input. A further aspect to address is the zero imputation. scRNA-seq data often have a significant amount of zeros. The problem of data zeros, especially structural zeros, is probably the thorniest and least-resolved issue in the log-ratio approach and has been called the “Achilles heel of compositional data analysis”. To apply CLR, these zeros must be transformed. While numerous strategies exist for this purpose, we have chosen, for the sake of simplicity, to add a minimal value 10^-4^ to each gene expression value, aiming to retain the originality of most values:

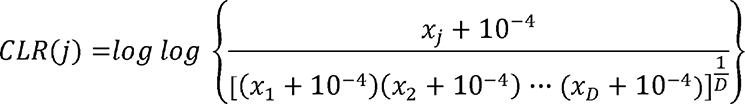

We performed an analysis on 10 out of 24 PBMC datasets (Table S1) to motivate and demonstrate the utility of the CLR transformation and balancing. For classification, the fine resolution dataset was used, it was split into 75% training and 25% testing sets. Lasso was employed as the classifier with 10-fold cross-validation for hyperparameter tuning. Evaluation metrics were training and testing error. For clustering, coarse-resolution dataset was used, the dimensionality was first reduced to 20 principal components before applying *k-*means with the number of clusters set to the known number of cell types. Evaluation metrics were Adjusted Rand Index (ARI) and Normalized Mutual Information (NMI). The same random seed was used across methods for reproducibility. All results were averaged over 10 datasets, as shown in Table 2, compared with raw data without any transformation, traditional log-normalization decreased testing error from 0.172 to 0.157, but did not decrease the training error for classification. For clustering, traditional log-normalization improved ARI from 0.448 to 0.529 and NMI from 0.550 to 0.602. CLR transformation largely improved performance for classification and clustering, reducing training error to 0.015 and testing error to 0.142 while increasing ARI to 0.527 and NMI to 0.604. Upon balancing the data that underwent CLR transformation, there was a notable improvement: the training error dropped to 0.0003 and the testing error decreased to 0.075, while both ARI and NMI saw increases to 0.724 and 0.798, respectively. These outcomes are highly encouraging, underscoring the effectiveness of CLR transformation coupled with data balancing in capturing the compositional nature of scRNA-seq data, thereby enhancing the performance of classification and clustering in subsequent analyses. For traditional log-normalization, denote *x_i_*(*i* ɛ {1, …, *N*}) as the vector of gene expression value for cell *i,* the library size is set to be 10^5^ the formula is:

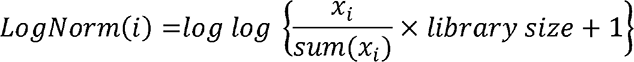

**Table 2.**
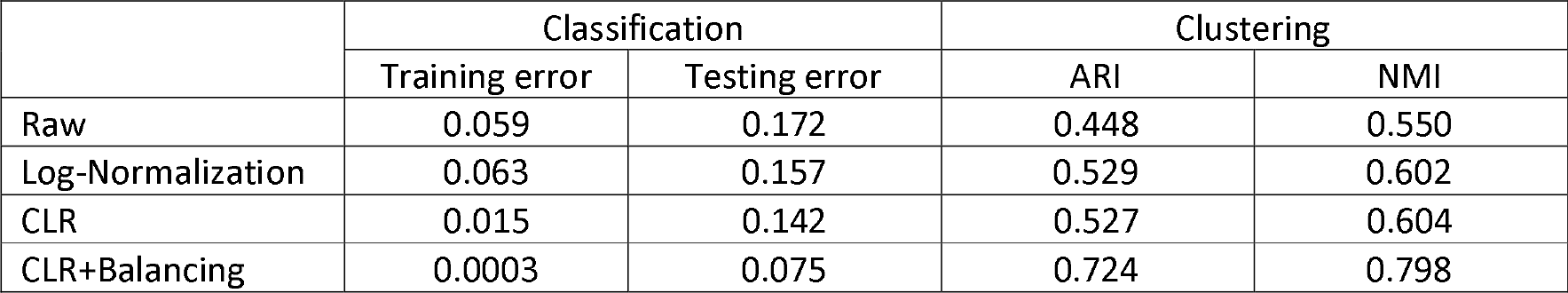
Motivation of CLR transformation based on classification and clustering performance.

When comparing log-normalization and centered log-ratio transformation, it’s evident that log-normalization is applied at the cell level, whereas CLR is targeted at the gene level. In compositional data analysis, genes represent the fundamental units of interest, rather than cells.

### 2.5 Machine learning classifiers and evaluation metrics

In our study, we employed 6 major categories of popular used supervised machine learning and deep learning classifiers evaluated for optimal compatibility with our proposed framework. Through conducting multi-classification and binary classification tasks on real world scRNA-seq datasets (in the results section), we aim to identify the most effective classifier compatible with our framework. A summary for each classification method are provided in Table 3. We evaluated 6 major classification methods: 1). Random Forest (RF); 2). Shrinkage methods (Lasso, ridge regression, elastic net with α = 0.5); 3). XGBoost; 4). *k-*Nearest Neighbors (*k-* NN); 5). Support Vector Machine (SVM); 6). Neural Networks (Multi-Layer Perceptron - MLP) *k*-NN was used for multi-class classification only, while SVM for binary classification only. Both used kernel techniques for enhanced performance in *k-*NN and SVM. The other classifiers were applied for both multi-class and binary cases. The key difference was the objective function - multinomial/softmax for multi-class and logistic for binary except for the non-model based *k*-NN method. We tailored preprocessing pipelines based on best practices for each algorithm. For shrinkage and neural networks methods, feature scaling handled varying magnitudes. For *k*-NN PCA was used for dimensionality reduction to improve high-dimensional performance. For RF and MLP, which do not inherently require PCA for dimensionality reduction, we still applied PCA as a pre processing step. The rationale was to limit memory usage and runtimes to feasible levels for model development on personal computers, aligning with our goal of enabling individual researchers without access to high-powered computing. Also, we found that the PCA transformation did not substantially impact model performance. For all methods using PCA for pre-processing, first 100 PCs were extracted as input features. For MLP, we evaluated 2, 3, and 4 layer architectures from simple to complex. All models were trained via 10-fold cross-validation for hyperparameter tuning. Default parameters were used otherwise in R.

**Table 3.**
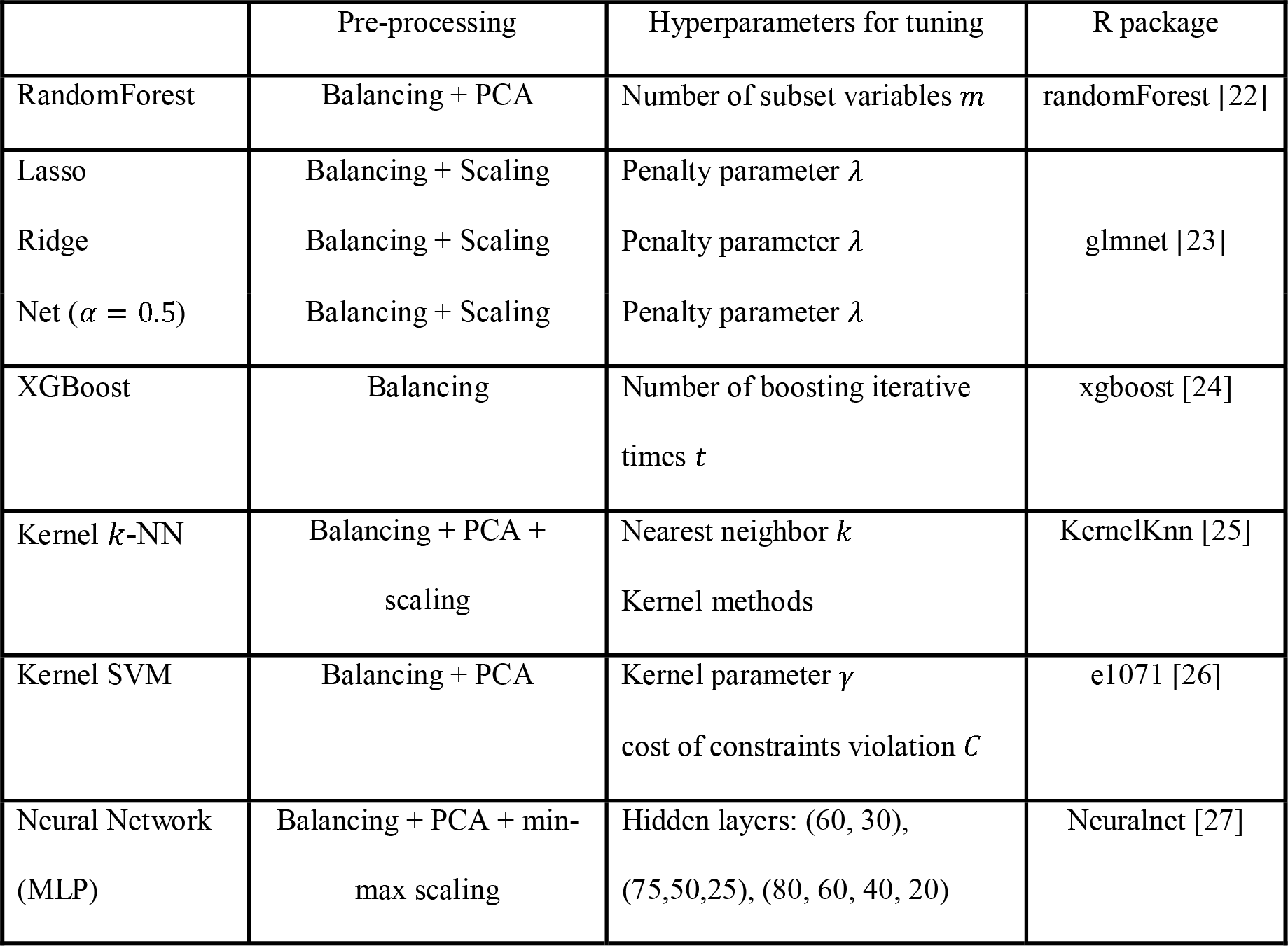
Summary of machine/deep learning classifier used in this study.

|  | Pre-processing | Hyperparameters for tuning | R package |
| --- | --- | --- | --- |
| RandomForest | Balancing + PCA | Number of subset variables $m$ | randomForest [22] |
| Lasso | Balancing + Scaling | Penalty parameter $\lambda$ | glmnet [23] |
| Ridge | Balancing + Scaling | Penalty parameter $\lambda$ | |
| Net ($\alpha = 0.5$) | Balancing + Scaling | Penalty parameter $\lambda$ | |
| XGBoost | Balancing | Number of boosting iterative times $t$ | xgboost [24] |
| Kernel $k$-NN | Balancing + PCA + scaling | Nearest neighbor $k$<br>Kernel methods | KernelKnn [25] |
| Kernel SVM | Balancing + PCA | Kernel parameter $\gamma$<br>cost of constraints violation $C$ | e1071 [26] |
| Neural Network (MLP) | Balancing + PCA + min-max scaling | Hidden layers: (60, 30), (75,50,25), (80, 60, 40, 20) | Neuralnet [27] |

In our study, we evaluated and compared our results using seven key classification evaluation metrics: 1) Accuracy, 2) Sensitivity, 3) Specificity, 4) Precision, 5) F-1 Score, 6) AUC (Area Under the Curve), and 7) AUC-PR (Area Under the Precision-Recall Curve). AUC-PR, similar to traditional AUC, uses precision and recall as its axes, making it more suitable for imbalanced datasets focused on optimizing the positive class. For binary classification, these metrics were calculated using a 2 × 2 confusion matrix. For multi-class classification, each *N* × *N* confusion matrix was broken down into *N* 2× *2* matrices, averaging the results of each metric. However, for multi-class tasks, we also calculated overall accuracy without splitting it into smaller matrices by simply dividing the diagonal sums by total sums.

## 3. Results

In scRNA-seq automated cell type annotation, the main goal is to assign each cell’s identity as accurately as possible. In supervised machine learning, this translates to minimizing classification error using our proposed framework. Before determining the best classifier, we validated that imbalance and compositional correction provides benefits under our framework in Section 3.2. Then, we evaluated both multi-class and binary classification to compare machine learning classifiers and identify the most compatible one, covered in Sections 3.3 and 3.4. Finally, Section 3.5 presents simulation studies exploring the framework’s ability to handle varying cell type proportions, demonstrating robustness. Before that, Section 3.1 shows some results on balanced data and related consequences. Overall, we comprehensively evaluated the proposed framework for scRNA-seq cell type annotation through multi-class and binary classification, validation of key components, and assessment across varying data distributions.

### 3.1 Data balancing results

Figure 2 shows UMAP visualizations of the PBMC training set before and after balancing. The top plots are the training coarse resolution dataset (Table S1) and the bottom plots are the training fine resolution dataset (Table S1). The left plots are before imbalance correction and the right plots are after correction. The coarse resolution data more clearly separates cell types, while fine resolution types like T cells overlap. Before correction, both resolutions are extremely imbalanced - some types like B cells (coarse) and MAIT cells (fine) are hard to discern in green/red on the left. After balancing, these minority types form tight clustered islands, visually confirming effective balancing. While coarse data more easily distinguishes cell types, the minority populations are well balanced after correction in both resolutions as evident by the emergent clusters.

**Figure 2:**
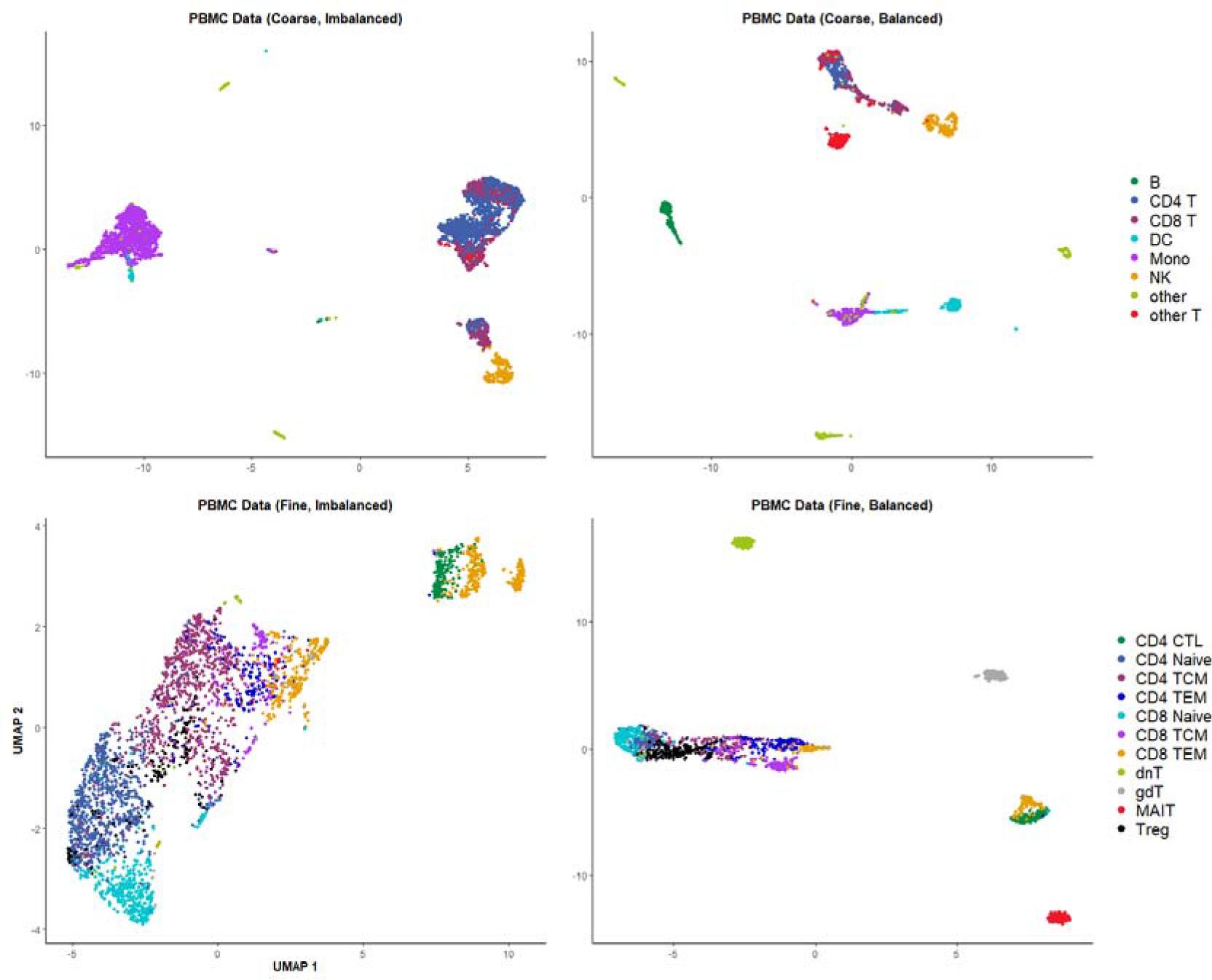
UMAP visualizations of PBMC training set before (left) and after (right) balancing.

Figure 3 shows the proportions of each cell type before (red bars) and after (blue bars) balancing for the PBMC coarse resolution (left) and fine resolution (right) training sets. Before balancing, proportions were extremely imbalanced in both resolutions. In the coarse data, CD4 T cells have ∼ 20× more samples than the minority B cells. After balancing, all cell types have equal 12.5% proportions. Similarly for the fine resolution data, balancing results in uniform proportions across types. The initially skewed distributions are transformed to well-balanced datasets. Correcting the class imbalance in this manner can help build more precise machine learning classifiers.

**Figure 3.**
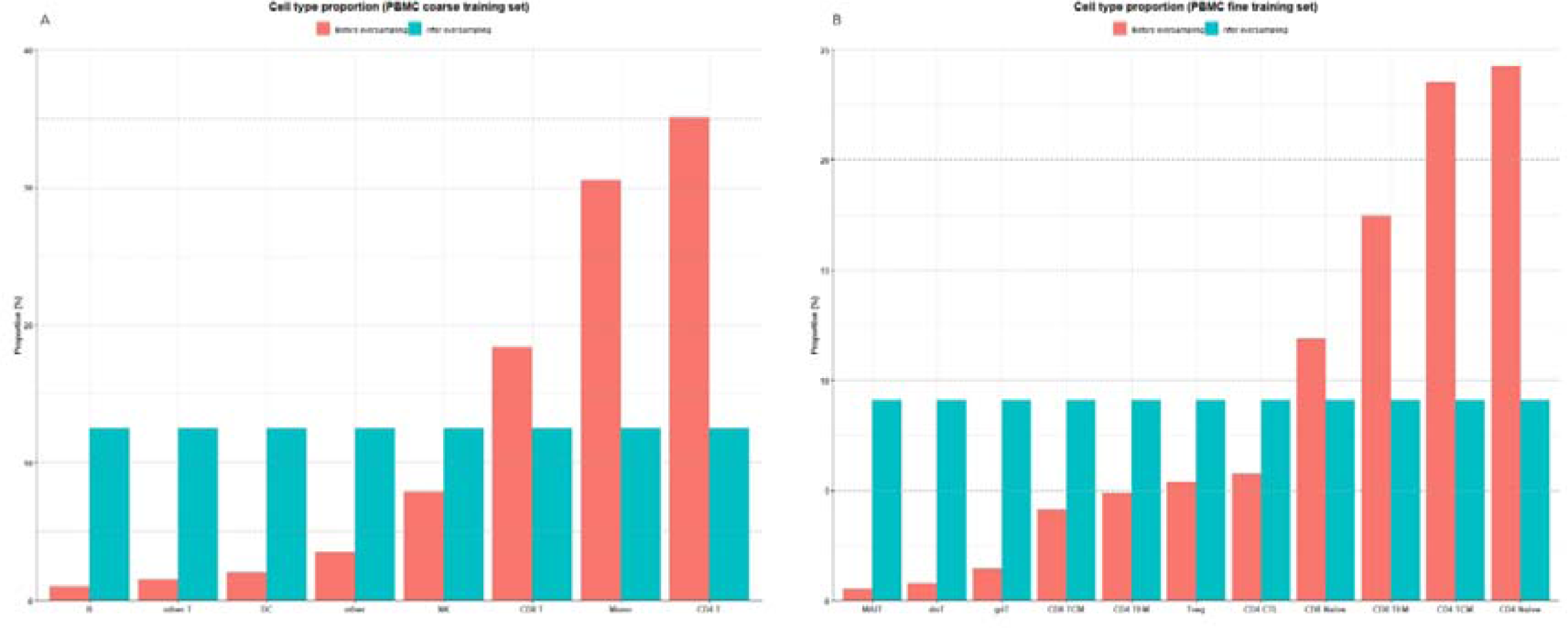
Proportions of cell types before (red) and after (blue) balancing of PBMC coarse (left) and fine (right) datasets.

Figure 3 shows the log expression density (*log*(*x* + 1)) for minority cell types in coarse (B cells) and fine (MAIT cells) resolutions. The blue density is the original minority cells, and red is the oversampled minority cells using our method. Our oversampling captures the expression profile well - each gene expression peak is simulated at similar levels as the original. We quantified this by the Kullback-Leibler (KL) divergence, which measures difference between probability distributions - smaller values indicate higher similarity. The KL divergence between original and oversampled distributions is small, indicating our oversampling method effectively models the scRNA-seq data. By replicating the expression density, our approach provides realistic synthetic minority samples for balancing.

As mentioned in Sections 2.2 and 2.3, to avoid losing valuable majority class information Figure 2 shows UMAP visualizations for the first layer coarse and fine resolutions. Figure S1 and Figure S2 display UMAP visualization of the other 4 of the 5 total layers after balancing. Imbalance correction works consistently across layers, with similar visualizations. Subjectively, the training sets appear suitable for building the machine learning classifiers. By repeating balancing, each layer retains all cell types with diversity maintained via different undersampling. This layered approach prevents discarding potentially useful majority samples. The consistent balanced distributions validate generating multiple complementary training sets.

### 3.2 Imbalance and compositional correction comparison

As mentioned in section 2.2, our proposed framework involves two key steps - imbalance correction and compositional correction. Before identifying the best machine learning classifier compatible with our framework, we first compared four scenarios across two resolutions to validate that both imbalance and compositional correction are necessary. The four scenarios are: 1) training data without imbalance correction or compositional transformation (No/No); 2) training data without imbalance correction but with compositional transformation (No/Yes); 3) training data with imbalance correction but no compositional transformation (Yes/No); and 4) training data with both imbalance correction and compositional transformation. Table 4 displays the final results of four scenarios on two data resolutions using XGBoost for multi-class classification. Except for the first metric column, other columns are the average metrics when splitting the *N* × *N* confusion matrix (where *N* is total number of cell types) into eight 2 × 2 matrices. Moreover, since we have 23 testing datasets that need annotation, we took the average of all evaluation metrics over the 23 datasets to obtain the final evaluation results, shown as mean and standard deviation. Values were multiplied by 100 for convenience. For multi-class classification, overall accuracy is more important than average accuracy since testing data is still imbalanced. Majority classes dominate accuracy, especially in 2 × 2 matrices. To consolidate coarse and fine resolution results for easier comparison, we combined the results of coarse and fine resolutions using utility weights of 0.3 for coarse and 0.7 for fine resolution, since real-world applications often focus more on classifying fine subtypes. The split table is shown in Table S3.

**Table 4:**
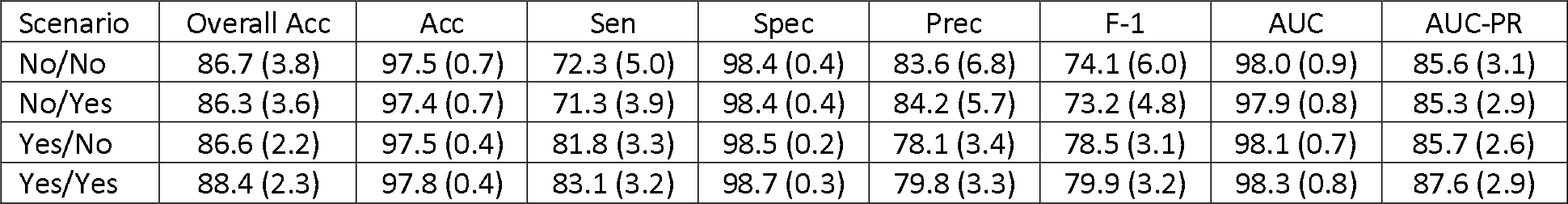
Evaluation metrics (multi-class) by different data processing scenarios on PBMC datasets (weighted average). (×100, Mean, SD)

| Scenario | Overall Acc | Acc | Sen | Spec | Prec | F-1 | AUC | AUC-PR |
| --- | --- | --- | --- | --- | --- | --- | --- | --- |
| No/No | 86.7 (3.8) | 97.5 (0.7) | 72.3 (5.0) | 98.4 (0.4) | 83.6 (6.8) | 74.1 (6.0) | 98.0 (0.9) | 85.6 (3.1) |
| No/Yes | 86.3 (3.6) | 97.4 (0.7) | 71.3 (3.9) | 98.4 (0.4) | 84.2 (5.7) | 73.2 (4.8) | 97.9 (0.8) | 85.3 (2.9) |
| Yes/No | 86.6 (2.2) | 97.5 (0.4) | 81.8 (3.3) | 98.5 (0.2) | 78.1 (3.4) | 78.5 (3.1) | 98.1 (0.7) | 85.7 (2.6) |
| Yes/Yes | 88.4 (2.3) | 97.8 (0.4) | 83.1 (3.2) | 98.7 (0.3) | 79.8 (3.3) | 79.9 (3.2) | 98.3 (0.8) | 87.6 (2.9) |

Clearly, imbalance correction increased sensitivity but decreases precision, while compositional transformation increased precision but decreased power. This occurred because models trained on imbalanced data can be biased toward majority classes, reducing detection power. Models lacking compositional transformation may not reflect real data relationships, hindering precision. Our proposed framework balanced sensitivity and precision, achieving the highest F1-score and AUC-PR on both coarse and fine resolution PBMC datasets. Except for precision, our framework achieved optimal performance across metrics. Although not the highest, precision only dropped slightly due to compositional transformation. Figure 4 visualizes these results in a forest plot, clearly demonstrating the optimal performance of our proposed framework with both imbalance correction and compositional transformation.

**Figure 4.**
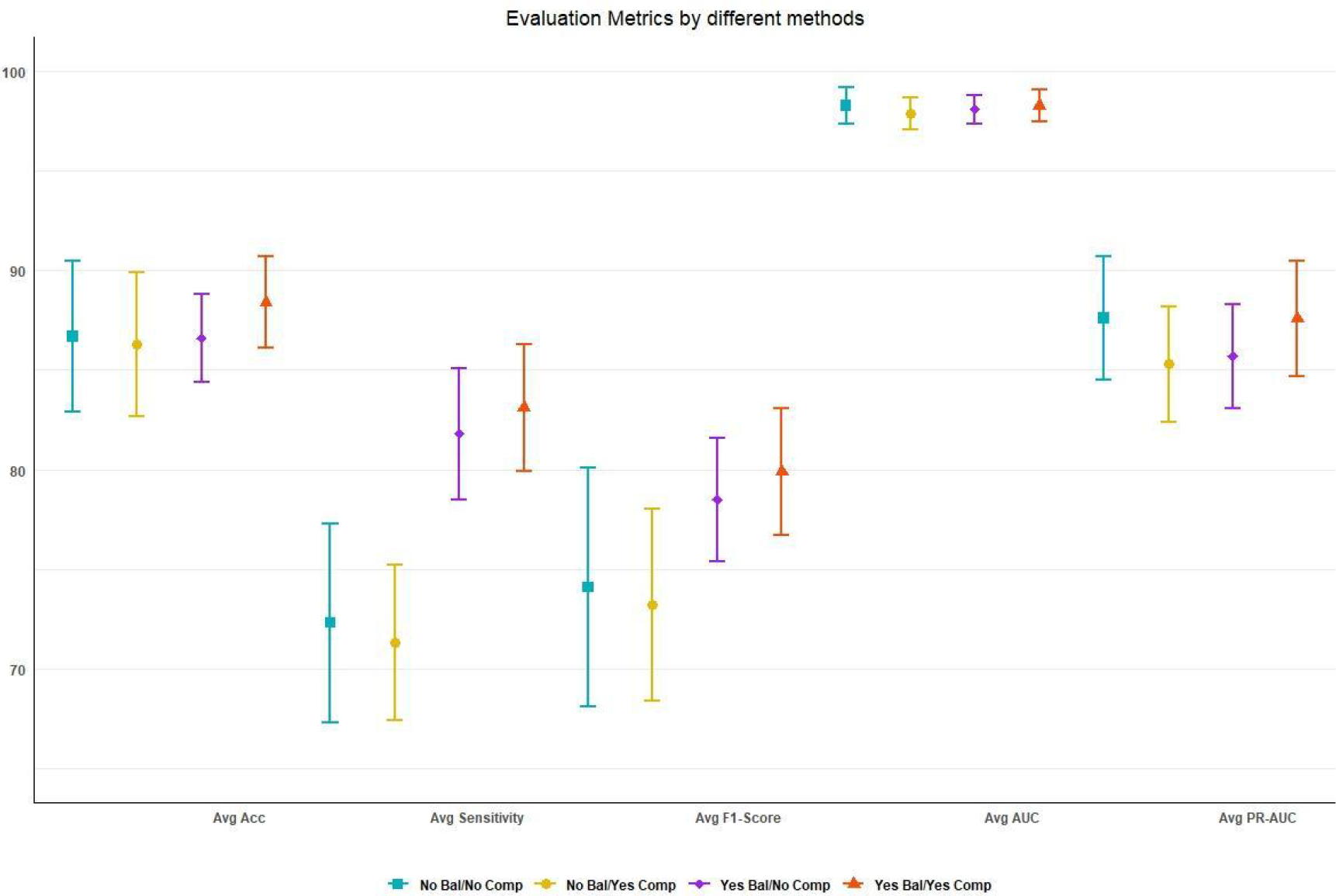
Forest plot for different data processing scenarios comparison.

### 3.3 Multi-class classification

Table S4 shows the evaluation metrics by different classifiers on PBMC coarse resolution datasets. The form of Table S4 is the same as Table 4 and Table S3. XGBoost had the best performance on 7 of 8 metrics, except sensitivity where elastic net with α =.0.5was best. Generally, XGBoost was superior based on these metrics for PBMC coarse data, with elastic net second. Notably, XGBoost had much higher precision and F1-score versus other lasso, ridge, and elastic net with α = 0.5 use shrinkage, performance differed. Lasso and classifiers, indicating high precision when predicting target cells as true labels. Although elastic net outperformed ridge regression since penalizing coefficients to 0 may be more beneficial given overfitting, where genes have correlation in scRNA-seq data. Additionally, deep learning MLP performed poorly, with 2 hidden layers best but still below non-deep methods. Hidden layers impacted results, but MLP was not optimal for this task.

Similarly, Table S5 shows evaluation metrics by classifier on PBMC fine resolution data. Overall, mean values were lower and standard deviations higher compared to coarse results. This aligns with expectations, as distinguishing fine cell subtypes within T cells is more challenging. Due to these intra-class similarities, performance metrics degrade and become more variable. XGBoost remained the top performer, with even larger advantages in precision and F1-score versus other classifiers. Notably, *k*-NN seemed to fail for the fine data when it succeeded for coarse resolution. The highly similar T cell subtypes likely embed closely together, hindering *k*--NN’s ability to distinguish them. Thus, the intrinsic challenges of fine classification become apparent, but XGBoost still emerged as the top method. Additionally, for fine resolution data MLP with 3 hidden layers performed best, unlike coarse resolution where 2 hidden layers were optimal. This highlights that different architectures suit different datasets, posing challenges for identifying the best neural network design when encountering new scRNA-seq data. While 2 hidden layers worked well for coarse classification, the complexities of fine subtype distinction benefited more from 3 hidden layers. Determining appropriate model complexity and architecture is a key consideration for effective application of MLP to varying scRNA-seq tasks.

Table 5 shows the final multi-class comparison using the same utility weights function in Section 3.2, clearly demonstrating XGBoost’s superiority and large advantages over other classifiers. As before, elastic net α = 0.5 ranked second. While we only showed α = 0.5 as an example, any α ɛ (0,1) could be used, with performance surpassing ridge regression and similar to lasso, indicating the benefits of regularizing to address overfitting. We cannot conclude the other classifiers are inherently poor, only that they are less compatible with our framework and scRNA-seq data. Nonetheless, XGBoost emerged as optimal for multi-class cell type annotation within our proposed ensemble balanced compositional framework. Figure 5 visualizes the results from Table 5, with each bar showing the mean of one evaluation metric for each classifier. Error bars indicate standard deviations, and the stars above the bars indicate significance levels from ANOVA tests, all with p-values < 0.001. This shows the classifier performance differences are statistically significant. The 4 key multi-class metrics are displayed. XGBoost’s superiority as the top performer is clearly visualized, consistent with the table. The figure summarizes the aggregated coarse and fine resolution results, highlighting XGBoost’s advantages across metrics with the smallest errors bars, demonstrating it as the most stable high-performing method.

**Figure 5.**
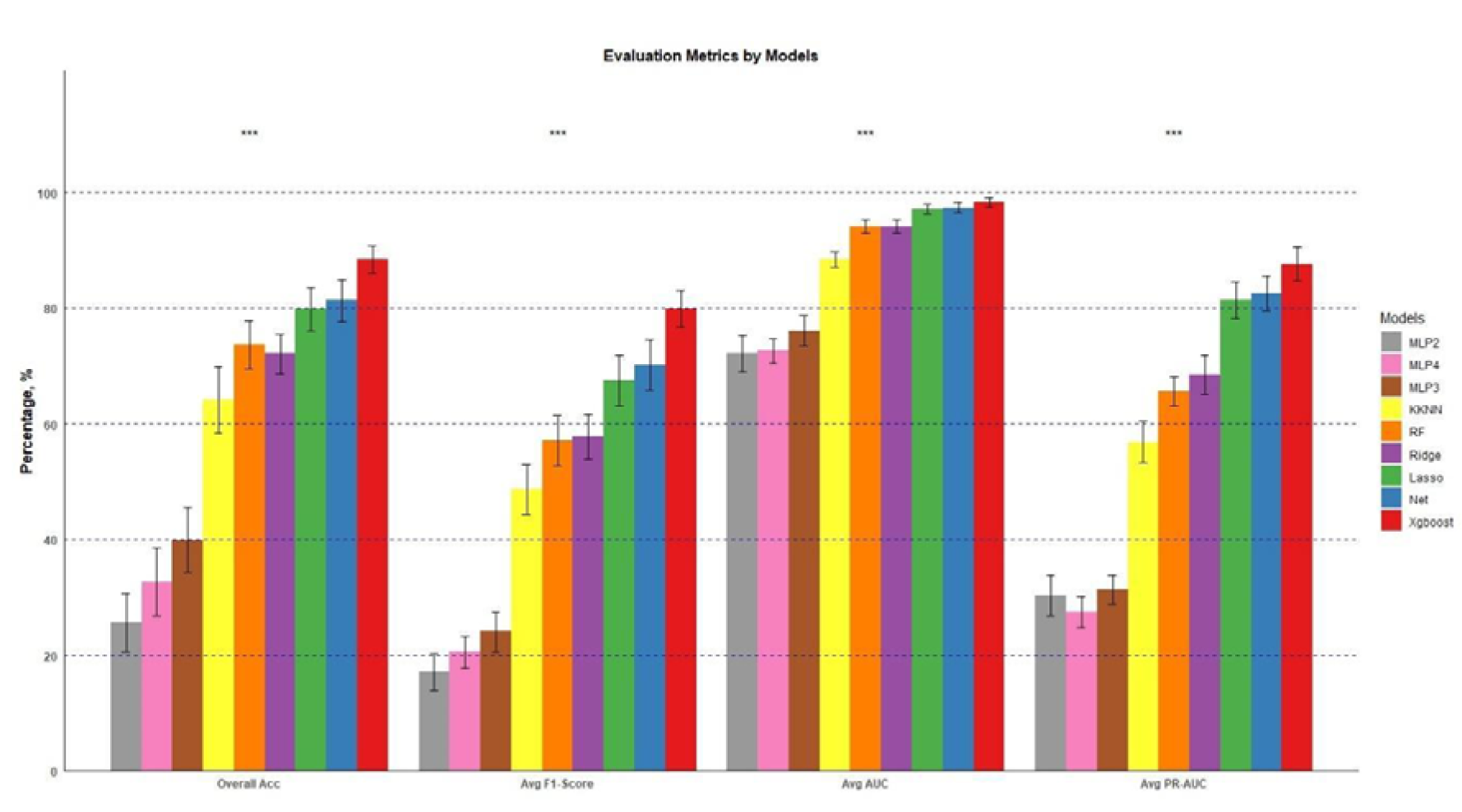
Barplot of evaluation metrics (multi-class) by different classifiers on PBMC datasets (weighted average).

**Table 5:** Evaluation metrics (multi-class) by different classifiers on PBMC datasets (weighted average). (×100, Mean, SD)

| Classifier | Overall Acc | Acc | Sen | Spec | Prec | F-1 | AUC | AUC-PR |
| --- | --- | --- | --- | --- | --- | --- | --- | --- |
| RF | 73.7 (4.2) | 94.9 (0.8) | 61.1 (3.2) | 97.0 (0.5) | 70.4 (4.7) | 57.1 (4.4) | 94.1 (1.1) | 65.7 (2.5) |
| Lasso | 79.8 (3.8) | 96.1 (0.7) | 81.0 (3.2) | 97.9 (0.4) | 64.2 (4.0) | 67.5 (4.3) | 97.1 (0.9) | 81.4 (3.2) |
| Ridge | 72.1 (3.4) | 94.7 (0.6) | 64.8 (3.2) | 97.0 (0.3) | 66.3 (4.0) | 57.8 (3.9) | 94.1 (1.1) | 68.5 (3.3) |
| Net ($\alpha = 0.5$) | 81.3 (3.6) | 96.4 (0.7) | 82.1 (3.0) | 98.0 (0.4) | 66.9 (4.1) | 70.2 (4.3) | 97.3 (0.9) | 82.5 (3.0) |
| XGBoost | 88.4 (2.3) | 97.8 (0.4) | 83.1 (3.2) | 98.7 (0.3) | 79.8 (3.3) | 79.9 (3.2) | 98.3 (0.8) | 87.6 (2.9) |
| Kernel $k$-NN | 64.2 (5.7) | 93.1 (1.1) | 54.0 (3.3) | 96.0 (0.6) | 63.5 (3.4) | 48.7 (4.3) | 88.4 (1.4) | 56.8 (3.6) |
| MLP (2 layers) | 25.7 (5.0) | 85.2 (1.1) | 23.8 (3.3) | 91.7 (0.5) | 38.0 (4.6) | 17.1 (3.2) | 72.2 (3.2) | 30.3 (3.5) |
| MLP (3 layers) | 39.9 (5.6) | 87.8 (1.2) | 31.8 (3.8) | 92.9 (0.6) | 35.0 (3.8) | 24.1 (3.4) | 76.1 (2.6) | 31.4 (2.5) |
| MLP (4 layers) | 32.7 (5.8) | 86.3 (1.2) | 27.2 (2.7) | 92.3 (0.6) | 29.8 (4.1) | 20.6 (2.8) | 72.6 (2.1) | 27.5 (2.7) |

Figure 6 displays performance of our framework with XGBoost on a same-donor coarse resolution testing set - donor 1 batch 2 (Table S1). Figure 6 (A) shows the UMAP visualization with true labels. Figure 6 (B) shows predictions from our framework with XGBoost classifier. The UMAPs appear very similar, with even some consistent labels between overlapped clusters. Figure 6 (C) is a confusion matrix heatmap comparing true and predicted labels. Diagonal values are accuracies in percentages. Off-diagonals show misclassification rates. Errors primarily occurred between T cell subtypes, understandable given their inherent similarities. Accuracy was strong for other cell types. Figure 6 (D) is a Sankey plot visualizing the annotation process, clearly reflecting the confusion matrix results. Overall, the consistent UMAPs, strong diagonal confusion matrix values, and informative Sankey plot validate our framework with XGBoost classifier’s ability to effectively predict cell types within the same donor.

**Figure 6.**
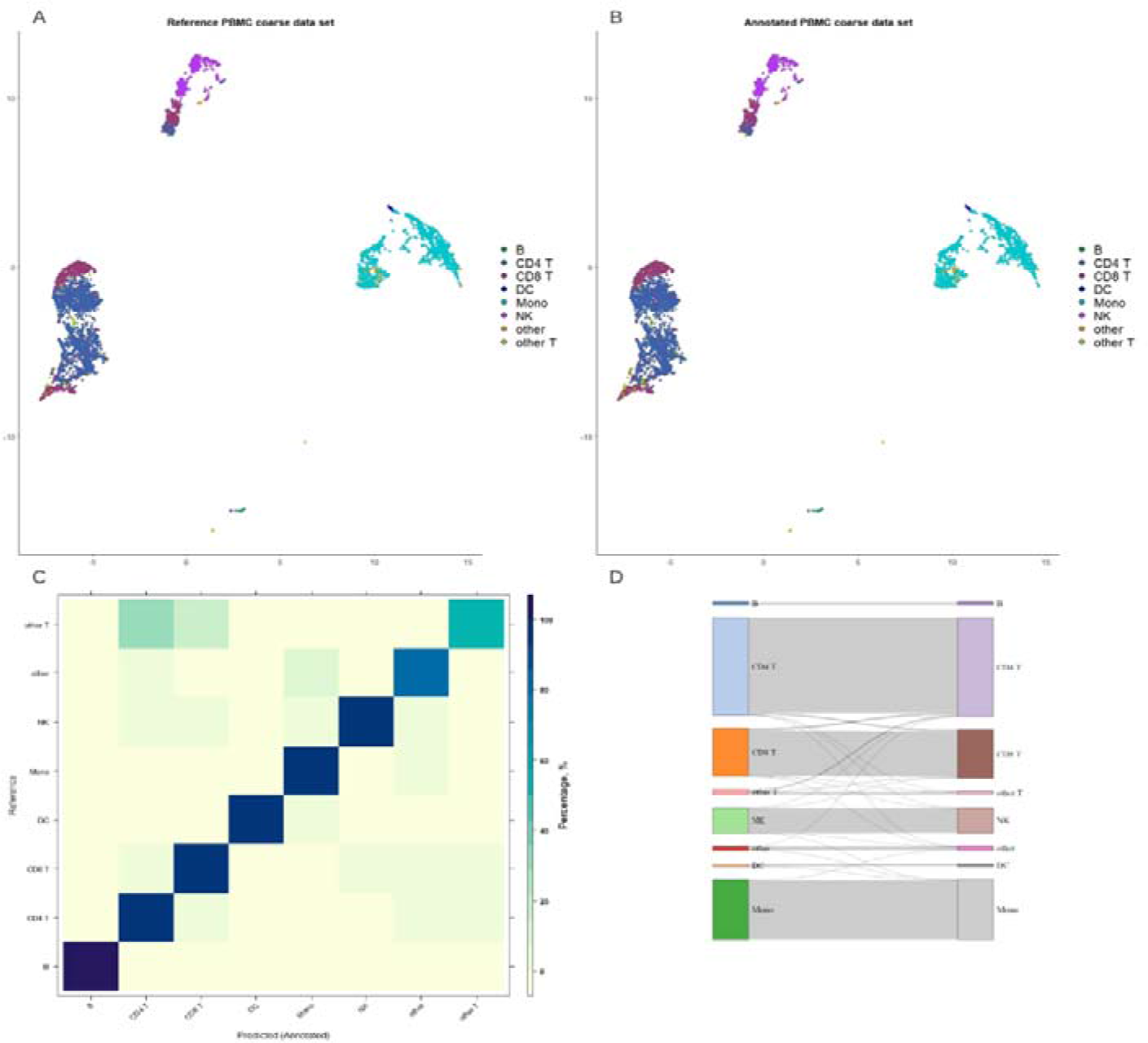
Annotation performance of ICCELF with XGBoost on a same-donor coarse resolution testing set. (A). UMAP visualization with true labels; (B). Annotated labels on the same dataset from the trained classifier; (C). Confusion matrix heatmap comparing true and predicted labels; (D). Sankey plot visualizing the annotation process.

Figure 7 displays performance of our framework with XGBoost on a same-donor fine resolution testing set - donor 1 batch 2 (Table S1). The interpretations of parts (A)-(D) of Figure 7 are analogous to those of Figure 6. The UMAP visualizations demonstrate that even for challenging to discriminate cell types in fine resolution datasets, our proposed framework implementing XGBoost as the classifier can achieve accurate predictions. Misclassifications predominantly transpired between CD4 TEM and CD4 TCM cells since they represent subtypes of CD4 T cells and therefore possess very similar properties. Additionally, some erroneous predictions surfaced between Treg cells and particular other T cell varieties.

**Figure 7.**
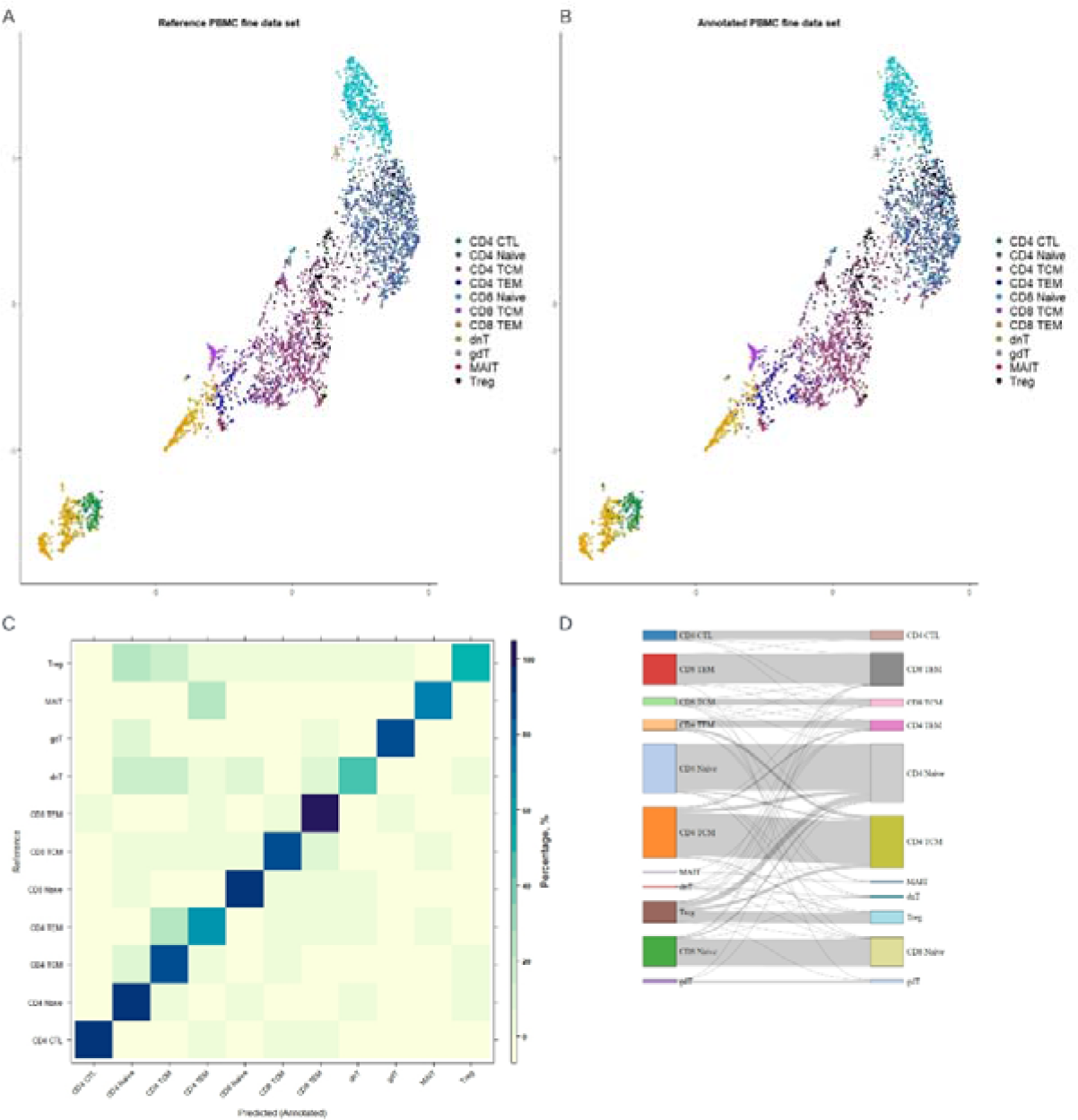
Annotation performance of ICCELF with XGBoost on a same-donor fine resolution testing set.

However, broadly speaking, the proposed framework integrating XGBoost as the classifier also attains sound performance on PBMC fine resolution datasets.

Figure 6 and Figure 7 exhibit the automated multi-class annotation outcomes for samples within the same donor, specifically donor 1. Conversely, Figure S2 and Figure S3 illustrate analogous results for PBMC coarse and fine resolutions but across discrete donors, namely donor 2 batch 1 (Table S1) as a representative example. We can discern that our proposed framework integrating XGBoost as the classifier can achieve robust performance when annotating samples across distinct donors.

### 3.4 Binary classification

In Section 2.3, we discussed the results of multi-class classification. Here in this section, we evaluated binary classifiers for the PBMC coarse resolution datasets. Since B cells comprise a small sample size for both the training and testing datasets in coarse resolution datasets, we considered B cells as the minority cell type and the target cell type for binary classification versus other cell types. Similarly, we considered gdT cells versus other cell types for binary classification on the fine resolution datasets. Table S6 presents evaluation metrics for binary classification by different classifiers on the PBMC coarse resolution datasets, analogous to Table S4. However, there are some differences. First, overall accuracy was discarded, since for binary classification, the confusion matrix is a 2 × 2 matrix, the overall accuracy is actually accuracy. Also, we replaced kernel *k*-NN with kernel SVM for the binary classification task. The results demonstrated that all classifiers performed well, even the MLP which failed for multi-class classification. The MLP model with 2 hidden layers showed particularly good results for binary classification. Both XGBoost and the SVM exhibited the best performance, while lasso had lower precision relative to the other models. The strong performance across models with little discrepancy is likely because B cells have distinct characteristics that make them easy to distinguish from other coarse resolution cell types.

Similarly, Table S7 presents results for binary classification of gdT cells versus other cell types in the PBMC fine resolution datasets. We observed that XGBoost outperformed other classifiers on accuracy, precision, F1-score, and AUC. Meanwhile, several classifiers that successfully made binary classifications on the coarse-resolution datasets failed on the fine-resolution dataset, including random forest and SVM. NA values indicate a denominator of 0 when calculating precision. These results illustrate that fine resolution data posed greater challenges for classification compared to coarse resolution data. The superior performance of XGBoost highlights its effectiveness at distinguishing gdT cells from other fine resolution cell types.

Table 6 shows the final binary classification comparison using the same utility weights function mentioned above. We observed similar results to the multi-class classification, with XGBoost as the top performer and elastic net with α = 0.5 as the second best. Figure 8 clearly visualizes these results in a bar plot. Figure 9 displays the ROC and PR-ROC plots, which correspond to the final AUC and AUC-PR values shown in Table 6. Specifically, we tracked the prediction values from each classifier for the coarse and fine resolution binary classification results. We then used the same utility weights to calculate the final AUC and precision-recall AUC, pooling the prediction values across the 23 testing datasets. This allows us to generate the final AUC and precision-recall AUC plots shown here in Figure 9.

**Figure 8.**
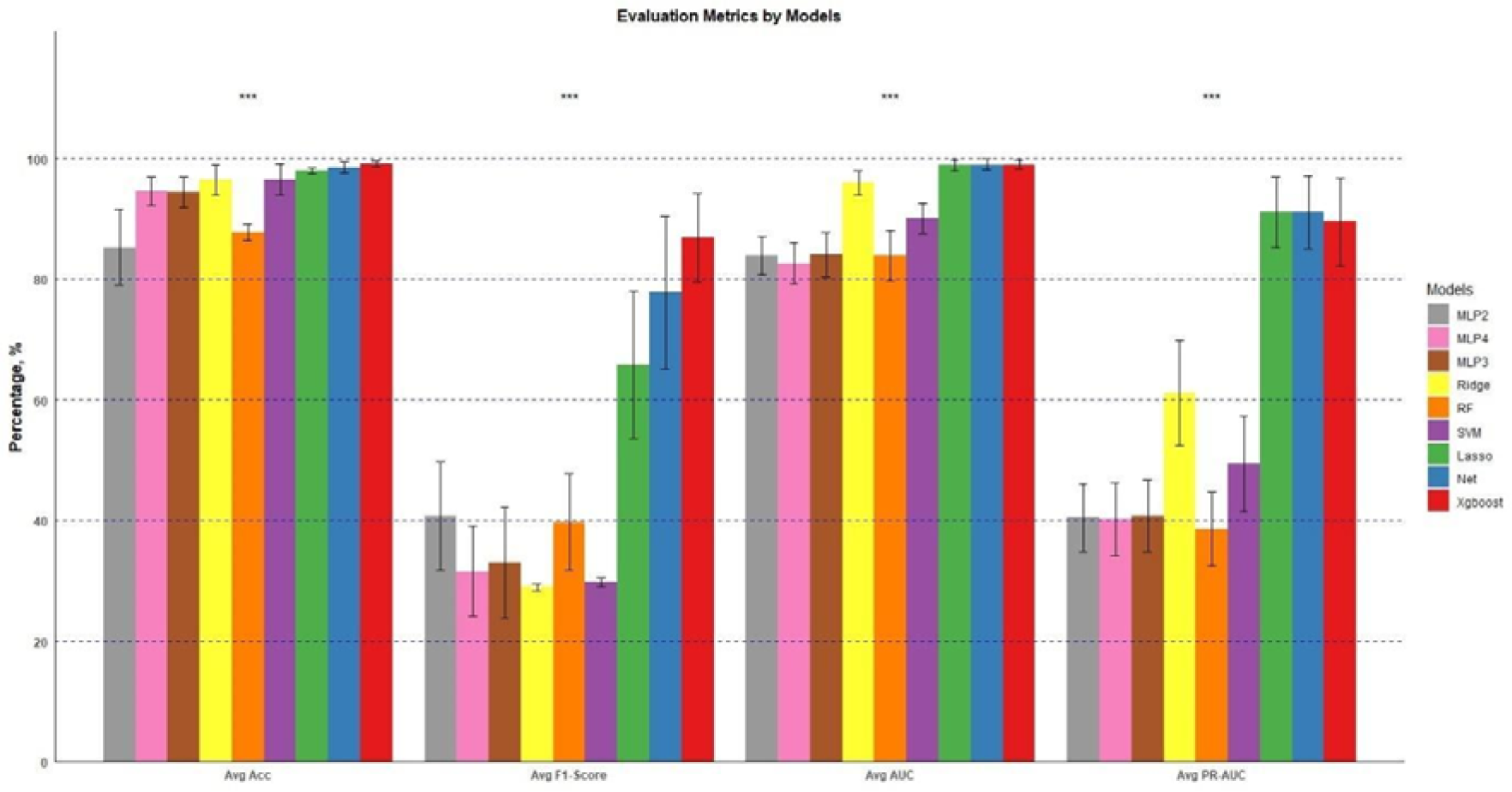
Barplot of evaluation metrics (binary) by different classifiers on PBMC datasets (weighted average)

**Figure 9.**
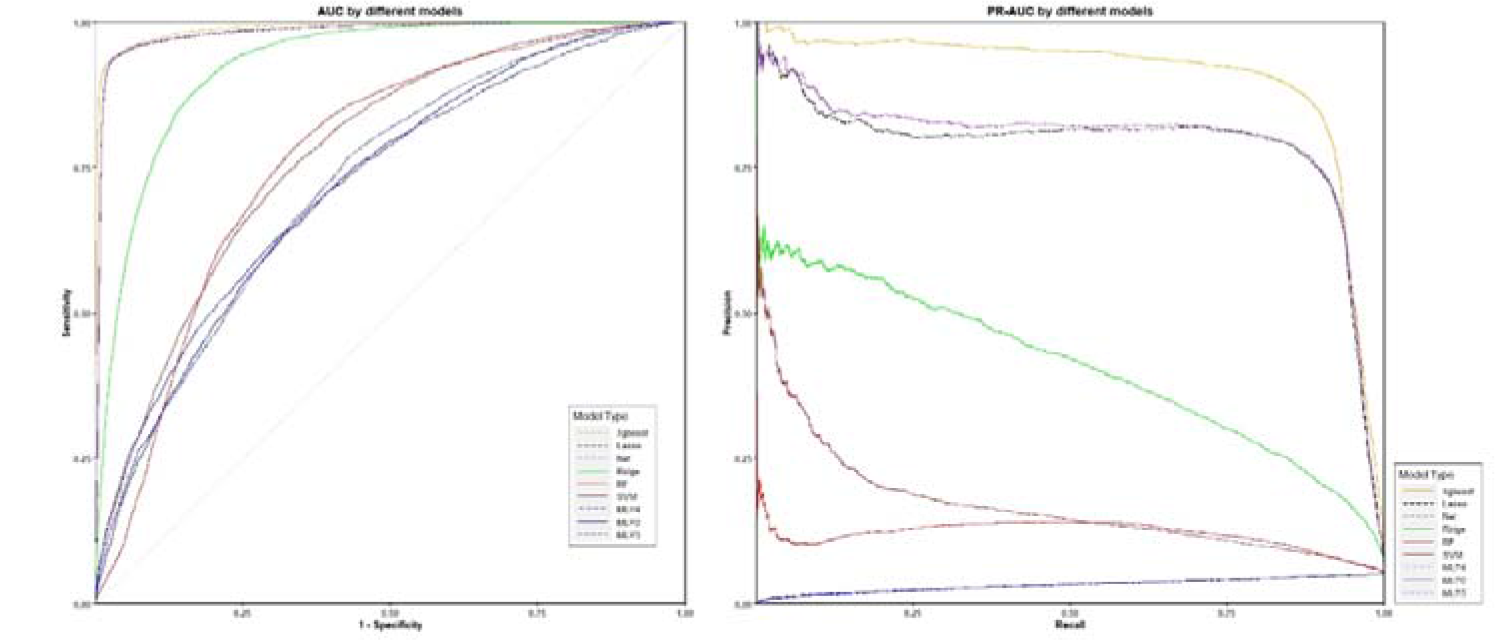
ROC (left) and PR-ROC (right) plots by different classifiers on PBMC datasets (weighted average).

**Table 6:** Evaluation metrics (binary) by different classifiers on PBMC datasets (weighted average).

| Classifier | Acc | Sen | Spec | Prec | F-1 | AUC | AUC-PR |
| --- | --- | --- | --- | --- | --- | --- | --- |
| RF | 87.7 (1.3) | 64.1 (10.1) | 89.1 (1.4) | 33.0 (6.9) | 39.8 (8.0) | 83.9 (4.2) | 38.6 (6.1) |
| Lasso | 97.9 (0.5) | 91.0 (4.9) | 98.4 (0.5) | 56.6 (17.1) | 65.8 (12.2) | 98.9 (0.9) | 91.1 (5.8) |
| Ridge | 96.5 (2.5) | 28.4 (1.1) | 100 (0) | NA | 29.0 (0.6) | 96.0 (2.0) | 61.2 (8.7) |
| Net ($\alpha = 0.5$) | 98.5 (0.9) | 88.3 (7.7) | 99.2 (0.5) | 73.7 (18.3) | 77.8 (12.6) | 99.0 (0.7) | 89.5 (7.2) |
| XGBoost | 99.1 (0.5) | 85.6 (7.8) | 99.6 (0.3) | 88.6 (7.9) | 86.9 (7.4) | 99.0 (0.7) | 89.5 (7.2) |
| Kernel SVM | 96.5 (2.6) | 29.8 (0.5) | 100 (0) | NA | 29.8 (0.8) | 90.1 (2.5) | 49.4 (7.9) |
| MLP (2 layers) | 85.3 (6.3) | 63.0 (10.8) | 86.6 (7.2) | 39.4 (6.7) | 40.8 (9.0) | 83.9 (3.2) | 40.5 (5.6) |
| MLP (3 layers) | 94.5 (2.5) | 32.2 (9.1) | 97.4 (2.2) | 47.9 (9.2) | 33.0 (9.2) | 84.0 (3.7) | 40.8 (5.9) |
| MLP (4 layers) | 94.6 (2.3) | 29.1 (6.6) | 97.7 (1.7) | 46.2 (10.3) | 31.6 (7.4) | 82.6 (3.4) | 40.2 (6.0) |

For all the binary classification results shown above, we focused on one target cell type versus other cell types, for both coarse and fine resolution datasets. However, the fine resolution datasets contain 11 different cell types. Therefore, we can build 11 binary classifiers, with results shown in Table S8 for models using our proposed framework with XGBoost as the classifier on the 11 distinct cell types in the fine-resolution datasets. We observed that models built for different cell types exhibited varying classification performance. gdT cells were the most readily distinguished cell type, while classifiers for other cell types like CD4 TCM and CD4 TEM had lower sensitivity and F1-scores, suggesting weaker distinction power. This is likely because CD4 TCM cells are very similar to some other CD4 T cell subtypes, making them harder for the model to identify. Figure S5 and Figure S6 provide corresponding barplot and ROC and PR-ROC plots for visual comparison, the high standard deviation indicates that our framework with XGBoost as the classifier achieved differing classification performance when built on different cell types. Compared to the multi-class classification results in Table 4, direct use of XGBoost for multi-class classification appears to be an effective choice, without the need to build 11 separate models which is more time-consuming. In summary, for our proposed ICCELF framework, XGBoost emerged as the top performing classifier on real-world PBMC datasets for both multi-class and binary classification tasks when compared to other machine learning classifiers. This superiority is most prominent for PBMC fine-resolution datasets, as highlighted in Sections 3.3 and 3.4.

### 3.5 Simulation results

As shown in Section 2.1 Table 1, we designated 5 scenarios simulating datasets ranging from heavily imbalanced to moderately imbalanced to balanced. This allows assessment across different imbalance levels while keeping other factors constant. Table 7 presents the simulation results across the 5 scenarios, using XGBoost as classifier for multi-class classification, the results were averaged over 20 testing sets. As balance increases from Scenario 1 to 5, all evaluation metrics showed an increasing trend. This is expected, as more balanced data typically improves machine learning classifier performance. However, the increases are minor, under 1% from Scenario 1 to 5. The forest plot in Figure 10 further illustrates the minimal differences between scenarios, with all evaluation metrics very similar for each scenario. This suggests that even with extreme imbalance, our proposed framework with XGBoost achieves classification performance comparable to balanced data.

**Table 7.** Evaluation metrics (multi-class) by different imbalance level scenarios on simulation datasets. (×100, Mean, SD)

| Scenarios | Overall Acc | Acc | Sen | Spec | Prec | F-1 | AUC | AUC-PR |
| --- | --- | --- | --- | --- | --- | --- | --- | --- |
| Scenario 1 | 94.4 (2.8) | 98.9 (0.6) | 94.4 (2.8) | 99.4 (0.3) | 95.2 (2.2) | 94.3 (2.9) | 100 (0) | 99.6 (0.3) |
| Scenario 2 | 95.0 (2.8) | 99.0 (0.6) | 95.0 (2.8) | 99.4 (0.3) | 95.7 (2.2) | 95.0 (2.8) | 100 (0) | 99.7 (0.3) |
| Scenario 3 | 95.0 (2.7) | 99.0 (0.5) | 95.0 (2.7) | 99.4 (0.3) | 95.8 (2.0) | 95.0 (2.8) | 100 (0) | 99.8 (0.2) |
| Scenario 4 | 94.8 (2.8) | 99.0 (0.6) | 94.8 (2.8) | 99.4 (0.3) | 95.7 (2.1) | 94.7 (2.9) | 100 (0) | 99.8 (0.2) |
| Scenario 5 | 95.4 (2.7) | 99.1 (0.5) | 95.4 (2.7) | 99.5 (0.3) | 96.1 (2.1) | 95.4 (2.7) | 100 (0) | 99.8 (0.2) |

Mean, SD)

**Figure 10.**
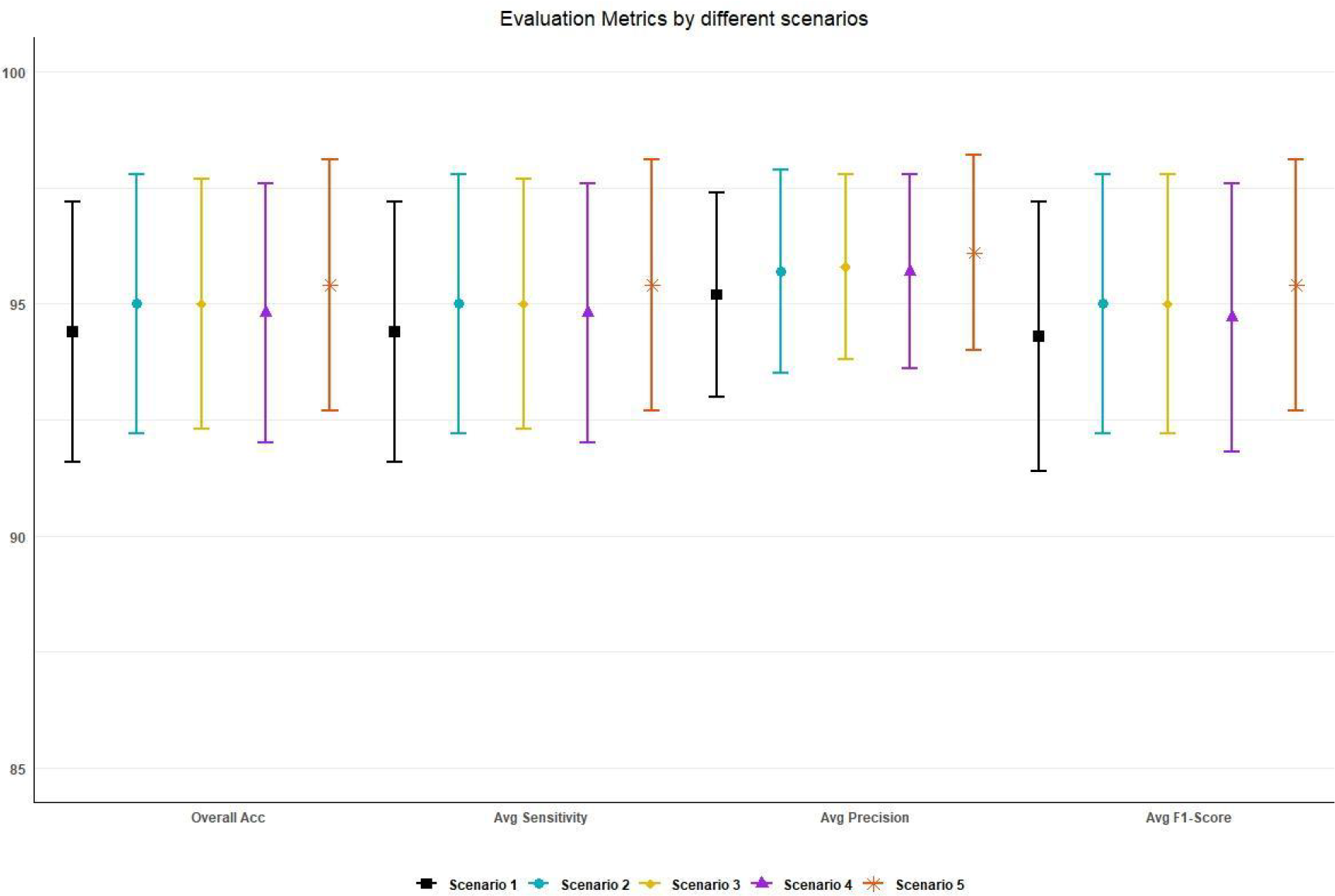
Forest plot for different data imbalance level scenarios comparison

## 4. Discussion

scRNA-seq produces high-dimensional gene expression profiles for individual cells. A key analysis task is annotating cell identity by assigning type labels. Manual annotation is time-consuming, subjective, and limits reproducibility. Automated supervised machine learning methods can enable rapid, scalable annotation. However, class imbalance and compositionality of scRNA-seq data pose challenges. Imbalanced cell type proportions bias machine learning classifiers, while compositionality violates assumptions of any standard multi-variate algorithms including most of machine learning classifiers.

This study introduces an Imbalance and Composition Corrected Ensemble Learning Framework to address these limitations. The framework integrates two key steps - imbalance correction via intelligent over/undersampling technique and compositional transformation using centered log-ratios. Multiple balanced and transformed training sets are generated to train an ensemble of classifiers for prediction for sake of obtaining robust results. This design balances sensitivity and precision while adhering to principles of compositional data analysis.

Comprehensive benchmarking of classification algorithms identified XGBoost as the optimal classifier compatible with ICCELF. For both multi-class and binary tasks, XGBoost significantly outperformed other methods like random forests, support vector machines, and neural networks on real PBMC datasets. Advantages were most prominent for fine resolution subtype classification, highlighting capabilities on complex heterogeneous cell populations. The superior performance of XGBoost can be attributed to its ensemble nature combining beneficial aspects of several machine learning approaches. As a tree-based method, it shares strengths of random forests like robustness to noise. By penalizing and shrinking coefficients like lasso and ridge regression, XGBoost handles feature collinearity. Its additive boosting provides advantages for sequential learning on structured data. Our framework generates layered synthetic training sets by combining real scRNA-seq data with oversampled minority classes. This structure is well-suited for XGBoost’s boosting approach. The additive training process enables effectively learning from both real and synthetic samples. In contrast, deep neural networks did not perform as well on our framework’s synthetic data structure. However, this does not imply neural networks are fundamentally ineffective. Rather, they may not be optimized for this particular dataset configuration. Additionally, XGBoost does not require extensive pre-processing like dimensionality reduction or feature scaling. It can be directly applied to the raw scRNA-seq data. This simplifies the workflow. XGBoost also exhibits computational efficiency.

While generally effective, our framework implementing XGBoost for binary classification of fine resolution cell subtypes exhibited limitations in certain cases. For intrinsically similar subtypes like CD4 TCM and CD4 TEM cells, the binary classification performance declined with lower sensitivity and F1-scores. This occurred because their inherent similarities made it difficult for the model to reliably distinguish them in a binary setting. However, multi-class classification using the same framework and XGBoost classifier performed well, accurately annotating amongst all fine subtypes simultaneously. Therefore, we recommend directly applying our proposed framework with XGBoost for multi-class classification when the goal is annotating an entire heterogeneous scRNA-seq dataset. Constructing multiple binary classifiers may not be necessary and could encounter limitations for closely related cell varieties. The multi-class approach can effectively categorize cells amongst all subtypes together. Binary classification may be more suitable for targeted detection or exploration of a specific putative novel cell type versus all other cells. But for routine annotation of known subtypes, multi-class classification integrates well with our imbalance and composition corrected ensemble framework.

We also validated that both imbalance correction and compositional transformation are essential components of the framework. Without these steps, metrics like sensitivity and precision declined due to biases and violated assumptions. The full proposed workflow with both corrections achieved highest performance by balancing precision and sensitivity. Testing on simulated scRNA-seq datasets with varying levels of class imbalance demonstrated consistent strong accuracy and F1-score. This highlights the framework’s robustness across diverse cell type distributions.

While promising, there are some limitations to note. First, it currently requires a single input training set which may not fully capture diverse cell types if limited in scope. Batch effects between training and deployment could also impact generalization. Integrating multiple datasets could provide more representative training data, potentially obviating the need for oversampling. However, integration risks introducing noise or bias if inaccurate methods are used. It also increases computational demands, conflicting with the goal of simplifying workflow for users. Second, benchmarking was conducted primarily on PBMC datasets as they are highly prevalent. However, numerous other tissue types can be profiled. Performance on additional sample sources beyond PBMCs requires further assessment. Third, while several major machine learning classifiers were evaluated, numerous other algorithms exist. Similarly, hyperparameter tuning was not exhaustive for each method. The conclusion that XGBoost is most compatible is reasonable based on results, but not definitive. More extensive comparisons on larger classifier sets could strengthen this claim. The accuracy of automated cell type annotation, which employs supervised machine learning classifiers, is significantly contingent on the precision of the annotated labels in the training set. Should the expert annotations in the training set be inaccurate, the resulting machine learning classifier built on this flawed foundation will inevitably produce unreliable annotations for new samples. In summary, limitations around training data representativeness, evaluation breadth, and result generalizability qualify the conclusions. But the framework shows promise as an accessible simplified solution. Expanding assessments and comparisons in future work can further optimize performance across diverse applications.

This work opens several future directions. The framework could be assessed on additional complex heterogeneous scRNA-seq datasets from various tissues and technologies. Comparisons to other state-of-the-art methods would better benchmark performance. The optimized classifiers could be deployed for production annotation or made publicly available. Biological insights from high-confidence automated mapping could reveal new cell states and trajectories. Overall, the proposed imbalance and composition corrected ensemble learning approach shows strong potential to accelerate reliable cell type annotation from scRNA-seq.

## Code Availability

The ICCELF function and all the R files for implementing the analysis can be accessed on the following GitHub repository: https://github.com/SaishiCui/ICCELF.

## Data Availability

Both the SPARSim simulation datasets and the real-world PBMC working datasets were generated using an R file available at https://github.com/SaishiCui/ICCELF.

Details about PBMC datasets can be found in Section 2. The original real-world PBMC datasets can be obtained from: https://satijalab.org/seurat/articles/multimodal_reference_mapping.html

## Author Contributions

SC and SN developed and designed the ICCELF framework. SC designed and wrote R programs. SC implemented and performed benchmarking analyses. SC wrote the manuscript. IFZ took responsibility and oversaw the mathematical and statistical theoretical foundation, IFZ and SN carried out revision of the manuscript.

## Acknowledgements.

The authors would like to acknowledge Dr. Sina Nassiri and Dr. Issa Zakeri for their mentorship and guidance, and for critical feedback and discussions. We thank the Department of Epidemiology and Biostatistics of Drexel University, Philadelphia PA, USA, for grants supporting this work.

## References

[1] . Clarke ZA, Andrews TS, Atif J, Pouyabahar D, Innes BT, MacParland SA, Bader GD. Tutorial: guidelines for annotating single-cell transcriptomic maps using automated and manual methods. Nature protocols. 2021 Jun;16(6):2749–64.

[2] . Pasquini G, Arias JE, Schäfer P, Busskamp V. Automated methods for cell type annotation on scRNA-seq data. Computational and Structural Biotechnology Journal. 2021 Jan 1;19:961–9.

[3] . Quinn TP, Erb I, Richardson MF, Crowley TM. Understanding sequencing data as compositions: an outlook and review. Bioinformatics. 2018 Aug 15;34(16):2870–8.

[4] . L. Lun AT, Bach K, Marioni JC. Pooling across cells to normalize single-cell RNA sequencing data with many zero counts. Genome biology. 2016 Dec;17:1–4.

[5] . Ahlmann-Eltze C, Huber W. Comparison of transformations for single-cell RNA-seq data. Nature Methods. 2023 May;20(5):665–72.

[6] . Hao Y, Hao S, Andersen-Nissen E, Mauck WM, Zheng S, Butler A, Lee MJ, Wilk AJ, Darby C, Zager M, Hoffman P. Integrated analysis of multimodal single-cell data. Cell. 2021 Jun 24;184(13):3573–87.

[7] . Baruzzo G, Patuzzi I, Di Camillo B. SPARSim single cell: a count data simulator for scRNA-seq data. Bioinformatics. 2020 Mar;36(5):1468–75.

[8] R Core Team. R: A language and environment for statistical computing [Internet]. Vienna, Austria: R Foundation for Statistical Computing; 2023. Version 4.3.1. Available from: https://www.R-project.org/

[9] . Crowell HL, Morillo Leonardo SX, Soneson C, Robinson MD. The shaky foundations of simulating single-cell RNA sequencing data. Genome Biology. 2023 Dec;24(1):1–9.

[10] . Cao Y, Yang P, Yang JY. A benchmark study of simulation methods for single-cell RNA sequencing data. Nature communications. 2021 Nov 25;12(1):6911.

[11] . Chawla NV, Bowyer KW, Hall LO, Kegelmeyer WP. SMOTE: synthetic minority over-sampling technique. Journal of artificial intelligence research. 2002 Jun 1;16:321–57.

[12] . Casella G, Berger RL. Statistical inference. Cengage Learning; 2021 Jan 26.

[13] . Vu TN, Wills QF, Kalari KR, Niu N, Wang L, Rantalainen M, Pawitan Y. Beta-Poisson model for single-cell RNA-seq data analyses. Bioinformatics. 2016 Jul 15;32(14):2128–35.

[14] . Risso D, Perraudeau F, Gribkova S, Dudoit S, Vert JP. A general and flexible method for signal extraction from single-cell RNA-seq data. Nature communications. 2018 Jan 18;9(1):284.

[15] Aitchison J. Monographs on statistics and applied probability. The statistical analysis of compositional data. 1986.

[16] . Van den Boogaart KG, Tolosana-Delgado R. Analyzing compositional data with R. Berlin: Springer; 2013 Jun 29.

[17] . Van den Boogaart KG, Tolosana-Delgado R. “Compositions”: a unified R package to analyze compositional data. Computers & Geosciences. 2008 Apr 1;34(4):320–38.

[18] . Lovell D, Pawlowsky-Glahn V, Egozcue JJ, Marguerat S, Bähler J. Proportionality: a valid alternative to correlation for relative data. PLoS computational biology. 2015 Mar 16;11(3):e1004075.

[19] . Aitchison J, Barceló-Vidal C, Martín-Fernández JA, Pawlowsky-Glahn V. Logratio analysis and compositional distance. Mathematical geology. 2000 Apr;32:271–5.

[20] . Aitchison J. The statistical analysis of compositional data. Journal of the Royal Statistical Society: Series B (Methodological). 1982 Jan;44(2):139–60.

[21] . Greenacre M, Grunsky E, Bacon-Shone J, Erb I, Quinn T. Aitchison’s compositional data analysis 40 years on: A reappraisal. Statistical Science. 2023 Jan;1(1):1–25.

[22] . Liaw A, Wiener M. Classification and regression by randomForest. R news. 2002 Dec 3;2(3):18–22.

[23] . Friedman J, Hastie T, Tibshirani R. Regularization paths for generalized linear models via coordinate descent. Journal of statistical software. 2010;33(1):1.

[24] Chen T, He T, Benesty M, Khotilovich V, Tang Y, Cho H, Chen K, Mitchell R, Cano I, Zhou T. Xgboost: extreme gradient boosting. R package version 0.4–2. 2015 Aug 1;1(4):1-4.

[25] . Mouselimis L. KernelKnn: Kernel k Nearest Neighbors [Internet]. R package version 1.1.5. 2023 [cited 2023 Feb 26]. Available from: https://CRAN.R-project.org/package=KernelKnn

[26] . Meyer D, Dimitriadou E, Hornik K, Weingessel A, Leisch F, Chang CC, Lin CC. e1071: misc functions of the department of statistics, probability theory group (formerly: E1071), TU Wien. R package version. 2019;1(2).

[27] . Günther F, Fritsch S. Neuralnet: training of neural networks. R J.. 2010 Jun 1;2(1):30.

