## Supplementary Methods for "Imbalance and Composition Correction Ensemble Learning Framework (ICCELF): A novel framework for automated scRNA-seq cell type annotation"

**Appendix A**

**Theorem** Testing of Poisson variance.

$H_{0}$: The sample distribution follows Poisson distribution (Not overdispersed)

$H_{a}$: The sample distribution follows Negative Binomial distribution (Overdispersed)

Step 1. If sample ${\{X}_{1}, X_{2},\ldots,X_{n}\}$ are from Poisson distribution, $\bar{X}$, is the sample mean, we can estimate the parameter of Poisson distribution using method of moments $\hat{\lambda}$ = $\bar{X}$.

Step 2. Generate $n$ i.i.d variables from Poisson($\hat{\lambda}$) denoted as $\{\tilde{X}_{1}, \tilde{X}_{2},\ldots,\tilde{X}_{n}\}$ repeating $B$ (typically a large number) times, then we have $B$ simulated samples $\{\tilde{X}_{1}, \tilde{X}_{2},\ldots,\tilde{X}_{n}{\}}_{1}^{B}$.

Step 3. We can get $B$ simulated variances from $\{\tilde{X}_{1}, \tilde{X}_{2},\ldots,\tilde{X}_{n}{\}}_{1}^{B}$, denoted as $S_{b}^{2}=\frac{1}{n-1}\sum_{i=1}^{n} (\tilde{X}_{n}-\bar{X}_{b})^{2}, b=1,\ldots,B$. Also, denote $S_{o}^{2}=\frac{1}{n-1}\sum_{i=1}^{n} (X_{i}-\bar{X})^{2}$ as the original sample variance.

Then the probability that ($P_{\hat{\lambda}}\left( S^{2}>S_{o}^{2} \right)$) the true sample distribution is overdispersed under Poisson distribution with parameter $\hat{\lambda}$ can be approximated by

$$\frac{1}{B}\sum_{b=1}^{B} I\left( S_{b}^{2}>S_{o}^{2} \right) \to P_{\hat{\lambda}}\left( S^{2}>S_{o}^{2} \right), \mathrm{as} B\to\infty$$

where $I(\cdot)$ is the indicator function, it equals 1 if $S_{b}^{2}>S_{o}^{2}$, otherwise 0.

Step 4. If the approximated $P_{\hat{\lambda}}\left( S^{2}>S_{o}^{2} \right)< \alpha$, $\alpha$ is the significance level, typically 0.05, then we reject the null hypothesis, and accept that the sample distribution is overdispersed under the estimated Poisson distribution.

**Appendix B**

**Table S1.** Summary statistics of 24 (8 donors, 3 batches) PBMC datasets.

| Datasets | | Coarse-resolution datasets | | Fine-resolution datasets | |
| --- | --- | --- | --- | --- | --- |
| Donor | Batch | Total number  of cells | Total number of detected genes | Total number  of cells | Total number of detected genes |
| Training | |  | |  | |
| 1 | 1 | 6,267 | 16,878 | 3,440 | 16,134 |
| Testing | |  | |  | |
| 1 | 2 | 5,708 | 16,423 | 3,481 | 15,828 |
| 1 | 3 | 5,626 | 16,473 | 3,011 | 15,734 |
| 2 | 1 | 5,568 | 16,362 | 3,434 | 1,5780 |
| 2 | 2 | 5,337 | 16,149 | 3,644 | 15,676 |
| 2 | 3 | 5,137 | 16,189 | 3,175 | 15,698 |
| 3 | 1 | 4,372 | 15,890 | 2,105 | 15,161 |
| 3 | 2 | 4,820 | 16,042 | 1,997 | 15,153 |
| 3 | 3 | 4,690 | 16,075 | 2,677 | 15,505 |
| 4 | 1 | 4,950 | 16,077 | 2,851 | 15,462 |
| 4 | 2 | 5,411 | 16,142 | 3,755 | 15,728 |
| 4 | 3 | 5,604 | 16,205 | 2,809 | 15,555 |
| 5 | 1 | 6,546 | 17,789 | 2,394 | 16,960 |
| 5 | 2 | 6,579 | 18,141 | 2,089 | 16,973 |
| 5 | 3 | 7,106 | 18,286 | 3,105 | 17,550 |
| 6 | 1 | 5,422 | 17,451 | 1,530 | 16,230 |
| 6 | 2 | 6,404 | 17,989 | 3,318 | 17,394 |
| 6 | 3 | 5,895 | 17,889 | 1,776 | 16,729 |
| 7 | 1 | 8,011 | 18,372 | 2,645 | 17,297 |
| 7 | 2 | 7,071 | 18,442 | 3,769 | 17,706 |
| 7 | 3 | 7,576 | 18,664 | 2,963 | 17,647 |
| 8 | 1 | 8,611 | 18,435 | 4,026 | 17,808 |
| 8 | 2 | 9,000 | 18,534 | 4,990 | 18,070 |
| 8 | 3 | 7,871 | 18,247 | 4,076 | 17,710 |

**Table S2.** Highly variable gene designation for simulated datasets

| Cell type | Driver genes designation | HVGs designation  (Drive gene excluded) | Number of HVGs (N, %)  (Driver gene included) |
| --- | --- | --- | --- |
| Type 1 | Gene 1-20 | Gene 201-400, 801-1000, 1401-1600 | 620 (4.1) |
| Type 2 | Gene 21-40 | Gene 401-600, 801-1000, 1201-1400, 1601-1800 | 820 (5.5) |
| Type 3 | Gene 41-60 | Gene 201-600, 1401-1600, 1801-2000 | 820 (5.5) |
| Type 4 | Gene 61-80 | Gene 401-600, 1001-1600, 1801-2000 | 1000 (6.7) |
| Type 5 | Gene 81-100 | Gene 201-400, 601-800, 1601-2000 | 820 (5.5) |
| Type 6 | Gene 101-120 | Gene 401-600, 1001-1200, 1401-1600, 1801-2000 | 820 (5.5) |
| Type 7 | Gene 121-140 | Gene 201-400, 601-800, 1201-1400 | 620 (4.1) |
| Type 8 | Gene 141-160 | Gene 601-800, 1001-1200, 1601-1800 | 620 (4.1) |
| Type 9 | Gene 161-180 | Gene 201-400, 1201-1400, 1601-2000 | 820 (5.5) |
| Type 10 | Gene 181-200 | Gene 801-1200, 1401-1800 | 820 (5.5) |

**Table S3.** Evaluation metrics (multi-class) by different data processing scenarios on PBMC datasets. (×100, Mean, SD)

| Scenario | Overall Acc | Acc | Sen | Spec | Prec | F-1 | AUC | AUC-PR |
| --- | --- | --- | --- | --- | --- | --- | --- | --- |
| No/No/Coarse | 94.7 (1.9) | 98.7 (0.5) | 85.5 (2.6) | 99.1 (0.3) | 95.2 (1.1) | 87.8 (2.5) | 99.0 (0.4) | 94.6 (1.5) |
| No/Yes/Coarse | 94.4 (1.7) | 98.6 (0.4) | 84.3 (1.7) | 99.1 (0.3) | 95.2 (0.9) | 86.7 (1.7) | 99.0 (0.4) | 94.7 (1.5) |
| Yes/No/Coarse | 94.3 (1.2) | 98.6 (0.3) | 90.5 (2.4) | 99.2 (0.2) | 88.9 (2.3) | 89.2 (2.2) | 99.0 (0.4) | 93.5 (2.0) |
| Yes/Yes/Coarse | 95.1 (1.4) | 98.8 (0.3) | 90.5 (2.4) | 99.3 (0.2) | 90.7 (2.1) | 90.2 (2.3) | 99.0 (0.4) | 93.9 (2.1) |
| No/No/Fine | 83.3 (5.0) | 97.0 (0.9) | 66.6 (7.0) | 98.1(0.5) | 78.7 (9.7) | 68.2 (8.5) | 97.6 (1.3) | 81.8 (4.5) |
| No/Yes/Fine | 82.8 (4.7) | 96.9 (0.9) | 65.7 (5.4) | 98.1 (0.5) | 79.5 (8.1) | 67.5 (6.7) | 97.4 (1.1) | 81.2 (4.1) |
| Yes/No/Fine | 83.3 (3.0) | 97.0 (0.6) | 78.1 (4.4) | 98.2 (0.3) | 73.5 (5.1) | 73.9 (4.4) | 97.6 (0.9) | 82.4 (3.5) |
| Yes/Yes/Fine | 85.5 (3.0) | 97.4 (0.6) | 80.0 (4.1) | 98.4 (0.3) | 75.2 (4.7) | 75.5 (4.2) | 98.0 (1.1) | 84.9 (3.8) |

**Table S4** Evaluation metrics (multi-class) by different classifiers on PBMC coarse resolution datasets (×100, Mean, SD).

| Classifier | Overall Acc | Acc | Sen | Spec | Prec | F-1 | AUC | AUC-PR |
| --- | --- | --- | --- | --- | --- | --- | --- | --- |
| RF | 84.9 (4.7) | 96.2 (1.2) | 77.9 (1.9) | 97.6 (0.9) | 80.1 (2.3) | 76.6 (3.0) | 96.2 (1.3) | 82.7 (1.3) |
| Lasso | 87.3 (2.4) | 96.8 (0.6) | 89.6 (2.1) | 98.3 (0.3) | 70.0 (4.4) | 75.2 (4.2) | 98.5 (0.6) | 92.2 (2.2) |
| Ridge | 88.6 (2.8) | 97.2 (0.7) | 85.2 (2.7) | 98.4 (0.4) | 77.7 (3.8) | 79.8 (3.3) | 97.6 (1.2) | 89.5 (2.7) |
| Net ($\alpha=0.5$) | 89.7 (1.9) | 97.4 (0.5) | 90.9 (2.2) | 98.6 (0.2) | 75.6 (4.5) | 80.5 (3.9) | 98.7 (0.5) | 92.9 (2.1) |
| XGBoost | 95.1 (1.4) | 98.8 (0.3) | 90.4 (2.1) | 99.3 (0.2) | 90.7 (2.1) | 90.2 (2.3) | 99.0 (0.4) | 93.9 (2.1) |
| Kernel $k$-NN | 79.5 (4.2) | 94.9 (1.0) | 73.0 (2.7) | 96.7 (0.8) | 81.5 (2.9) | 73.1 (2.6) | 94.5 (1.5) | 78.5 (1.8) |
| MLP (2 layers) | 36.6 (14.9) | 84.2 (3.7) | 39.5 (8.4) | 91.7 (1.8) | 56.5 (6.2) | 31.9 (8.1) | 81.6 (3.0) | 49.8 (6.1) |
| MLP (3 layers) | 35.5 (10.3) | 83.9 (2.6) | 36.4 (3.5) | 90.9 (1.2) | 42.8 (6.7) | 24.5 (4.9) | 76.9 (2.7) | 39.2 (4.1) |
| MLP (4 layers) | 30.3 (10.5) | 82.6 (2.6) | 35.0 (4.8) | 90.6 (1.0) | 38.1 (6.7) | 23.3 (4.1) | 76.9 (2.9) | 39.1 (3.7) |

| Classifier | Overall Acc | Acc | Sen | Spec | Prec | F-1 | AUC | AUC-PR |
| --- | --- | --- | --- | --- | --- | --- | --- | --- |
| RF | 68.9 (5.0) | 94.3 (0.9) | 53.8 (4.1) | 96.7 (0.5) | 66.3 (6.2) | 48.8 (5.5) | 93.2 (1.5) | 58.5 (3.6) |
| Lasso | 76.6 (5.0) | 95.7 (0.9) | 77.3 (4.4) | 97.7 (0.5) | 61.7 (4.9) | 64.2 (5.5) | 96.5 (1.2) | 76.8 (4.4) |
| Ridge | 65.0 (5.2) | 93.6 (1.0) | 56.0 (4.7) | 96.4 (0.5) | 61.5 (6.0) | 48.4 (5.7) | 92.6 (1.4) | 59.4 (4.5) |
| Net ($\alpha=0.5$) | 77.6 (4.8) | 95.9 (0.9) | 78.4 (4.1) | 97.8 (0.5) | 63.2 (5.1) | 65.8 (5.6) | 96.7 (1.2) | 78.0 (4.1) |
| XGBoost | 85.5 (3.0) | 97.4 (0.6) | 80.0 (4.1) | 98.4 (0.3) | 75.2 (4.7) | 75.5 (4.2) | 98.0 (1.1) | 84.9 (3.8) |
| Kernel $k$-NN | 57.6 (7.7) | 92.3 (1.4) | 45.8 (3.9) | 95.6 (0.7) | 55.8 (4.7) | 38.2 (5.3) | 85.7 (1.9) | 47.5 (5.0) |
| MLP (2 layers) | 20.9 (5.0) | 85.6 (0.9) | 17.1 (2.8) | 91.7 (0.3) | 30.2 (6.5) | 10.8 (2.8) | 68.2 (3.9) | 21.9 (3.5) |
| MLP (3 layers) | 41.9 (6.7) | 89.4 (1.2) | 28.8 (4.5) | 93.7 (0.6) | 31.6 (4.2) | 23.9 (3.9) | 75.7 (3.0) | 28.1 (2.8) |
| MLP (4 layers) | 33.7 (6.0) | 87.9 (1.1) | 23.8 (3.0) | 93.1 (0.5) | 26.2 (4.2) | 19.4 (3.2) | 71.0 (2.4) | 22.5 (3.0) |

**Table S5** Evaluation metrics (multi-class) by different classifiers on PBMC fine resolution datasets (×100, Mean, SD).

| Classifier | Acc | Sen | Spec | Prec | F-1 | AUC | AUC-PR |
| --- | --- | --- | --- | --- | --- | --- | --- |
| RF | 99.7 (0.2) | 99.7 (0.6) | 99.7 (0.2) | 78.8 (9.0) | 87.8 (5.6) | 100 (0) | 99.7 (0.3) |
| Lasso | 97.8 (0.4) | 99.7 (0.7) | 97.7 (0.4) | 33.0 (3.6) | 49.5 (4.1) | 100 (0) | 99.2 (1.0) |
| Ridge | 99.9 (0) | 94.6 (3.6) | 100 (0) | 99.2 (1.3) | 96.8 (1.9) | 100 (0) | 99.2 (1.0) |
| Net ($\alpha=0.5$) | 99.7 (0.1) | 99.8 (0.7) | 99.7 (0.1) | 81.0 (3.9) | 89.4 (2.4) | 100 (0) | 99.4 (0.9) |
| XGBoost | 100 (0) | 98.7 (1.3) | 100 (0) | 98.5 (1.8) | 98.6 (1.4) | 100 (0) | 99.8 (0.4) |
| Kernel SVM | 100 (0) | 99.0 (1.2) | 100 (0) | 98.6 (1.9) | 98.8 (1.2) | 100 (0) | 100 (0.2) |
| MLP (2 layers) | 99.9 (0.2) | 87.7 (16.6) | 100 (0) | 99.4 (2.2) | 92.2 (10.7) | 100 (0.1) | 99.5 (0.8) |
| MLP (3 layers) | 99.5 (0.3) | 54.7 (25.3) | 100 (0) | 99.7 (0.9) | 66.8 (24.6) | 99.6 (0.4) | 94.2 (5.0) |
| MLP (4 layers) | 99.6 (0.2) | 59.5 (22.7) | 100 (0) | 99.8 (0.6) | 71.8 (20.5) | 99.8 (0.3) | 97.4 (2.7) |

**Table S6**. Evaluation metrics (binary) by different classifiers on PBMC coarse resolution datasets (×100, Mean, SD)

| Classifier | Acc | Sen | Spec | Prec | F-1 | AUC | AUC-PR |
| --- | --- | --- | --- | --- | --- | --- | --- |
| RF | 82.6 (1.8) | 48.8 (14.4) | 84.5 (1.9) | 13.4 (9.1) | 19.2 (11.4) | 77.0 (6.0) | 12.4 (8.7) |
| Lasso | 98.0 (0.7) | 87.2 (7.1) | 98.6 (0.7) | 66.7 (24.9) | 72.8 (18.1) | 98.4 (1.3) | 87.6 (8.3) |
| Ridge | 95.0 (3.6) | 0 (0) | 100 (0) | NA | 0 (0) | 94.4 (2.9) | 44.8 (12.4) |
| Net ($\alpha=0.5$) | 97.9 (1.2) | 83.4 (11.0) | 98.9 (0.7) | 70.6 (26.0) | 72.8 (17.7) | 98.5 (1.3) | 87.5 (8.5) |
| XGBoost | 98.7 (0.7) | 79.9 (10.8) | 99.5 (0.4) | 84.3 (11.3) | 81.8 (10.3) | 98.6 (0.9) | 85.1 (10.2) |
| Kernel SVM | 95.0 (3.6) | 0.1 (0.5) | 100 (0) | NA | 0.2 (0.9) | 85.8 (3.6) | 27.8 (11.3) |
| MLP (2 layers) | 79.0 (8.9) | 52.4 (16.3) | 80.9 (10.3) | 13.8 (9.7) | 18.7 (10.1) | 77.0 (4.6) | 15.2 (7.9) |
| MLP (3 layers) | 92.3 (3.6) | 22.6 (12.8) | 96.2 (3.2) | 25.7 (13.2) | 18.6 (7.0) | 77.3 (5.3) | 17.9 (7.8) |
| MLP (4 layers) | 92.5 (3.3) | 16.0 (8.3) | 96.6 (2.4) | 23.3 (14.7) | 14.4 (7.7) | 75.2 (4.8) | 15.6 (8.5) |

**Table S7**. Evaluation metrics (binary) by different classifiers on PBMC fine resolution datasets (×100, Mean, SD)

| (vs. Other cells) | Acc | Sen | Spec | Prec | F-1 | AUC | AUC-PR |
| --- | --- | --- | --- | --- | --- | --- | --- |
| gdT | 98.7 (0.7) | 79.9 (10.8) | 99.5 (0.4) | 94.3 (11.3) | 81.8 (10.3) | 98.6 (0.9) | 85.1 (10.2) |
| CD4 CTL | 99.1 (1.2) | 68.0 (31.6) | 99.7 (0.2) | 59.1 (37.2) | 56.2 (31.2) | 98.6 (3.8) | 64.8 (33.2) |
| CD4 Naïve | 84.5 (4.6) | 35.5 (14.8) | 99.8 (0.2) | 98.5 (1.4) | 50.4 (16.0) | 98.5 (0.8) | 95.3 (2.0) |
| CD4 TCM | 83.8 (6.3) | 25.5 (6.7) | 99.6 (0.3) | 95.4 (2.6) | 39.7 (8.4) | 94.3 (1.8) | 83.2 (5.7) |
| CD4 TEM | 94.4 (2.3) | 11.6 (7.5) | 99.8 (0.1) | 77.6 (19.7) | 19.4 (11.4) | 95.0 (2.3) | 58.6 (8.2) |
| CD8 Naïve | 96.7 (1.4) | 77.9 (6.5) | 99.8 (0.3) | 98.7 (1.5) | 86.9 (4.2) | 99.2 (0.7) | 97.4 (2.2) |
| CD8 TCM | 97.2 (1.9) | 39.8 (15.9) | 99.8 (0.2) | 87.1 (8.8) | 52.4 (17.0) | 95.3 (2.5) | 71.4 (9.7) |
| CD8 TEM | 95.7 (1.9) | 73.8 (12.0) | 98.5 (2.0) | 92.4 (4.6) | 81.4 (7.3) | 98.4 (0.7) | 92.4 (4.0) |
| dnT | 99.7 (0.3) | 49.8 (28.4) | 100 (0) | 87.6 (22.3) | 59.7 (25.3) | 91.2 (10.7) | 61.7 (24.5) |
| MAIT | 96.9 (2.5) | 26.5 (11.0) | 99.9 (0.1) | 92.3 (8.6) | 39.9 (14.6) | 99.4 (0.4) | 82.3 (15.1) |
| Treg | 98.2 (1.0) | 45.6 (17.5) | 99.7 (0.4) | 90.3 (9.7) | 57.9 (15.4) | 98.5 (1.0) | 81.7 (7.3) |

**Table S8**. Evaluation metrics (binary) by different cell types on PBMC datasets (XGBoost)

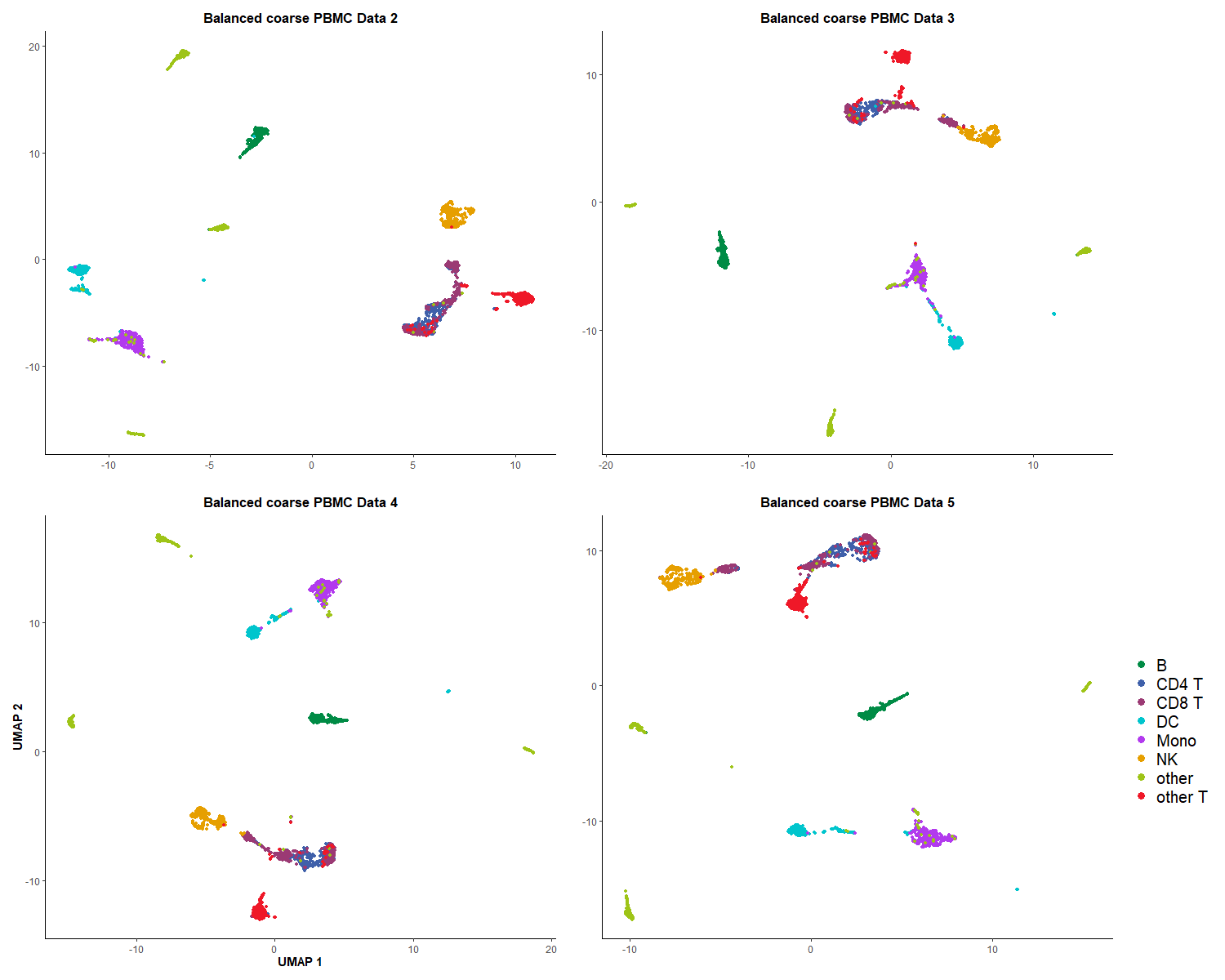

**Figure S1.** UMAP visualizations of PBMC coarse training set after balancing, the other 4 layers.

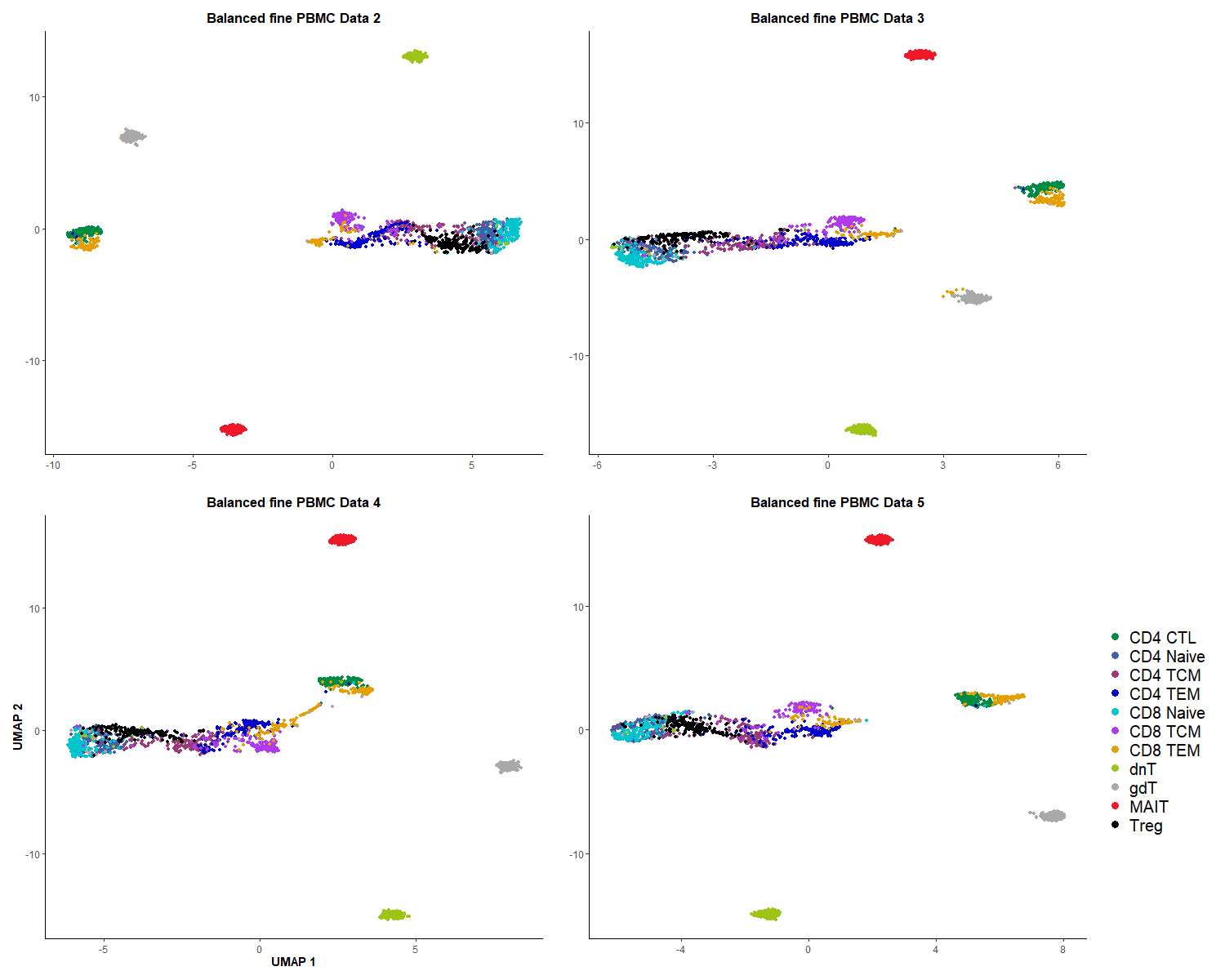

**Figure S2.** UMAP visualizations of PBMC fine training set after balancing, the other 4 layers.

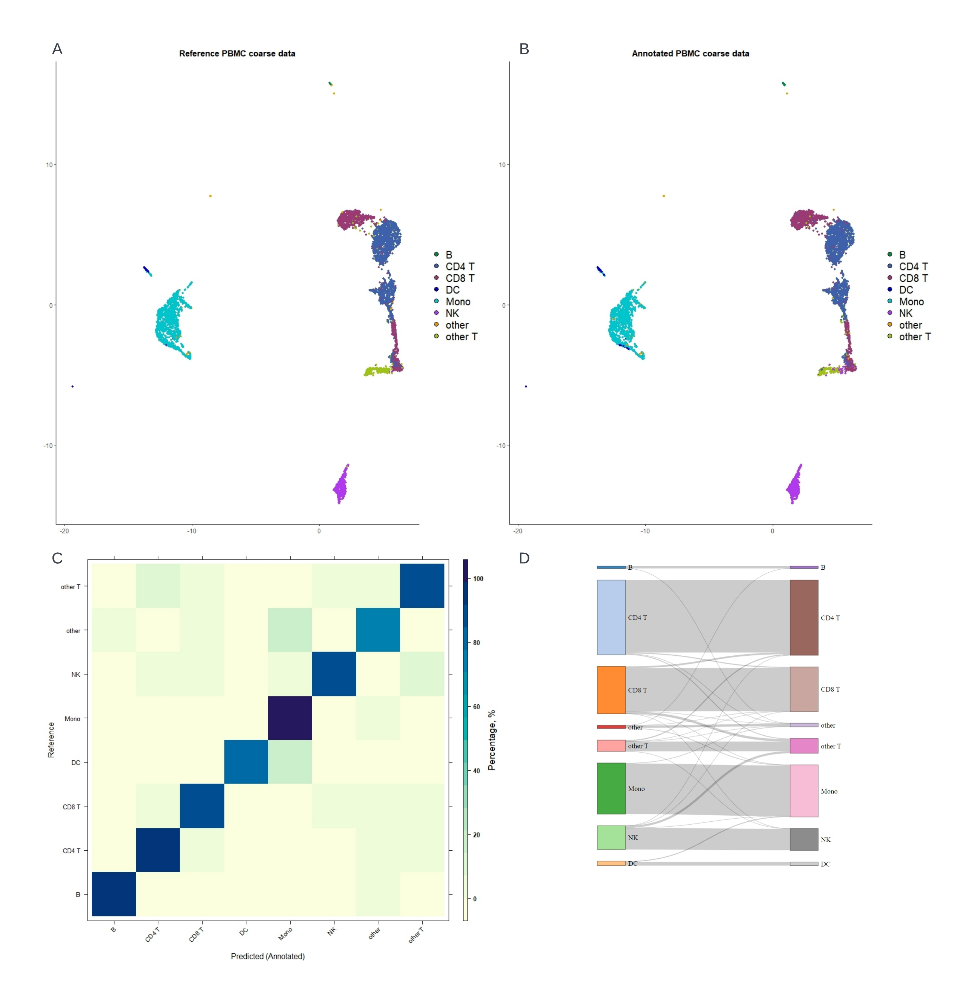

**Figure S3**. Annotation performance of ICCELF with XGBoost on a different-donor coarse resolution testing set.

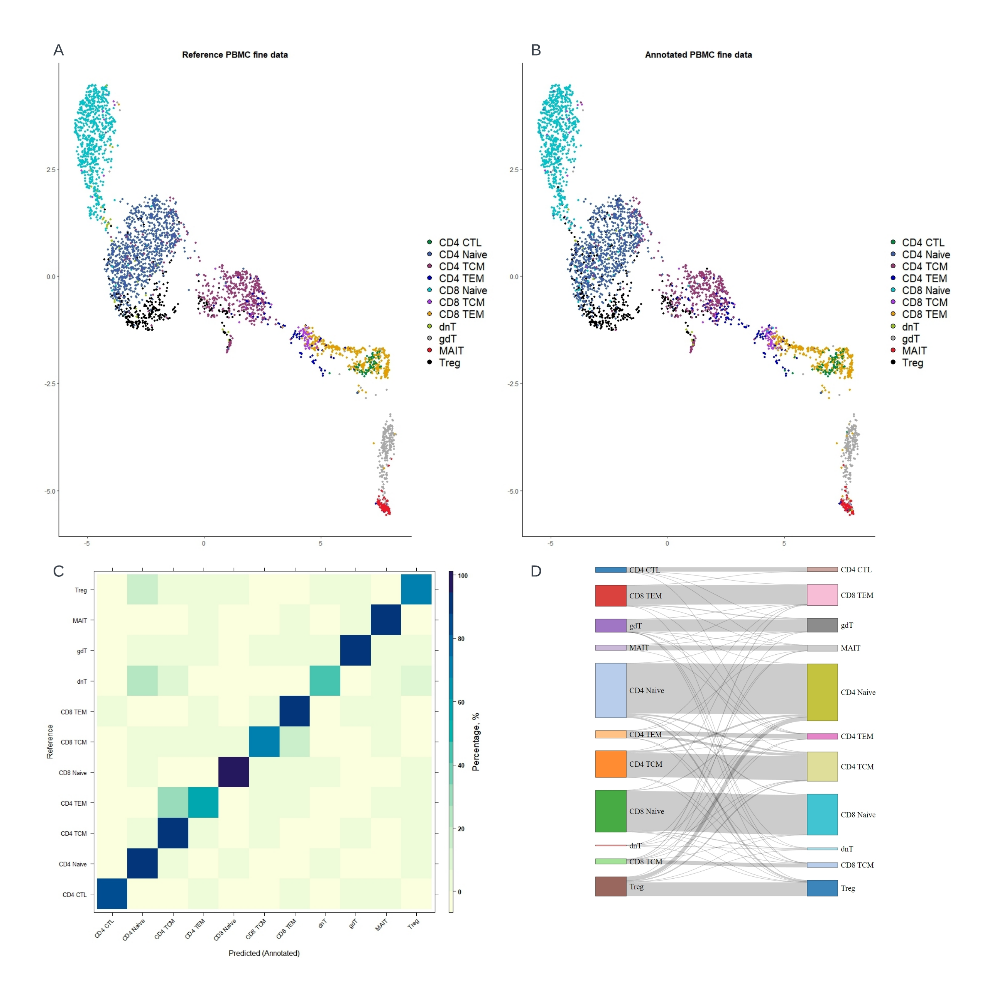

**Figure S4**. Annotation performance of ICCELF with XGBoost on a different-donor fine resolution testing set

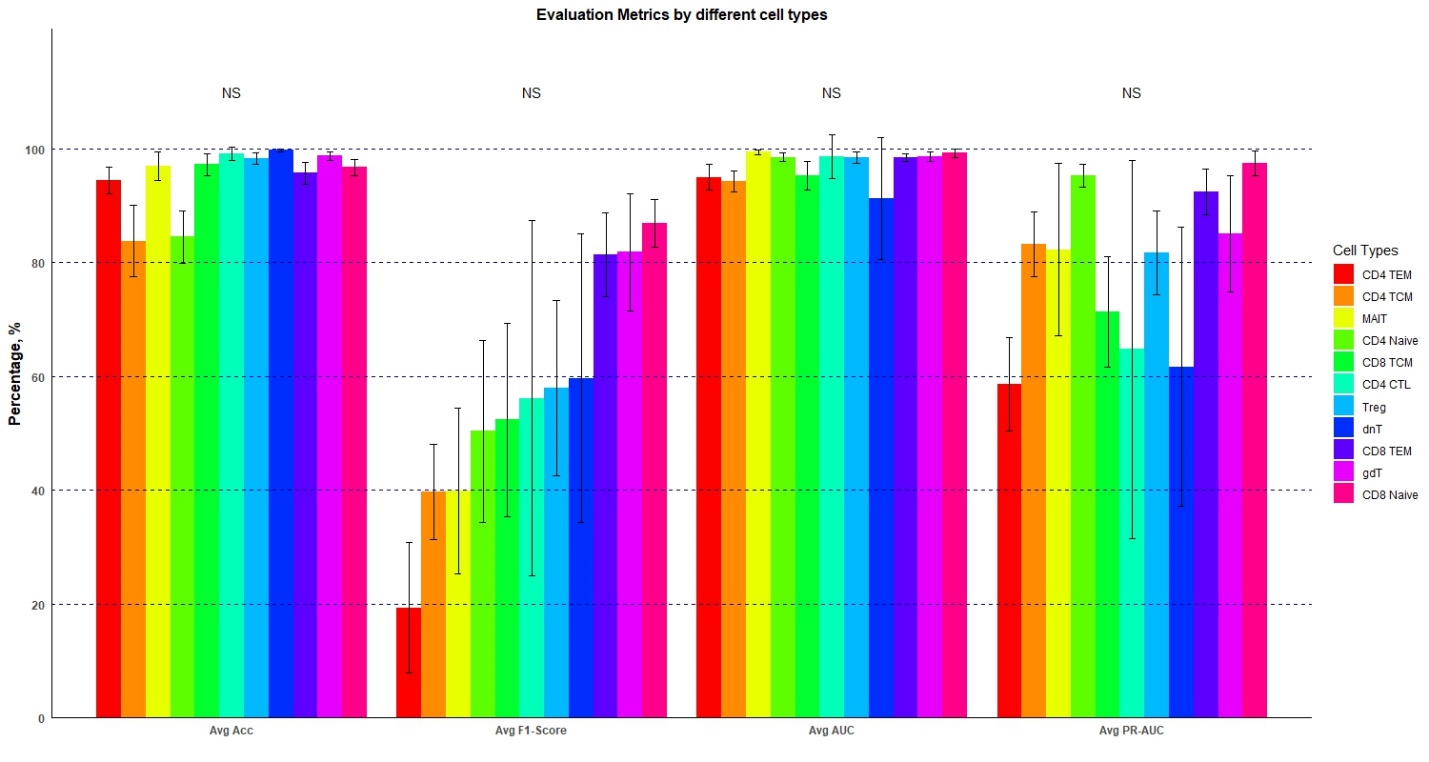

**Figure S5**. Barplot of evaluation metrics (binary) by different cell types on PBMC datasets (XGBoost)

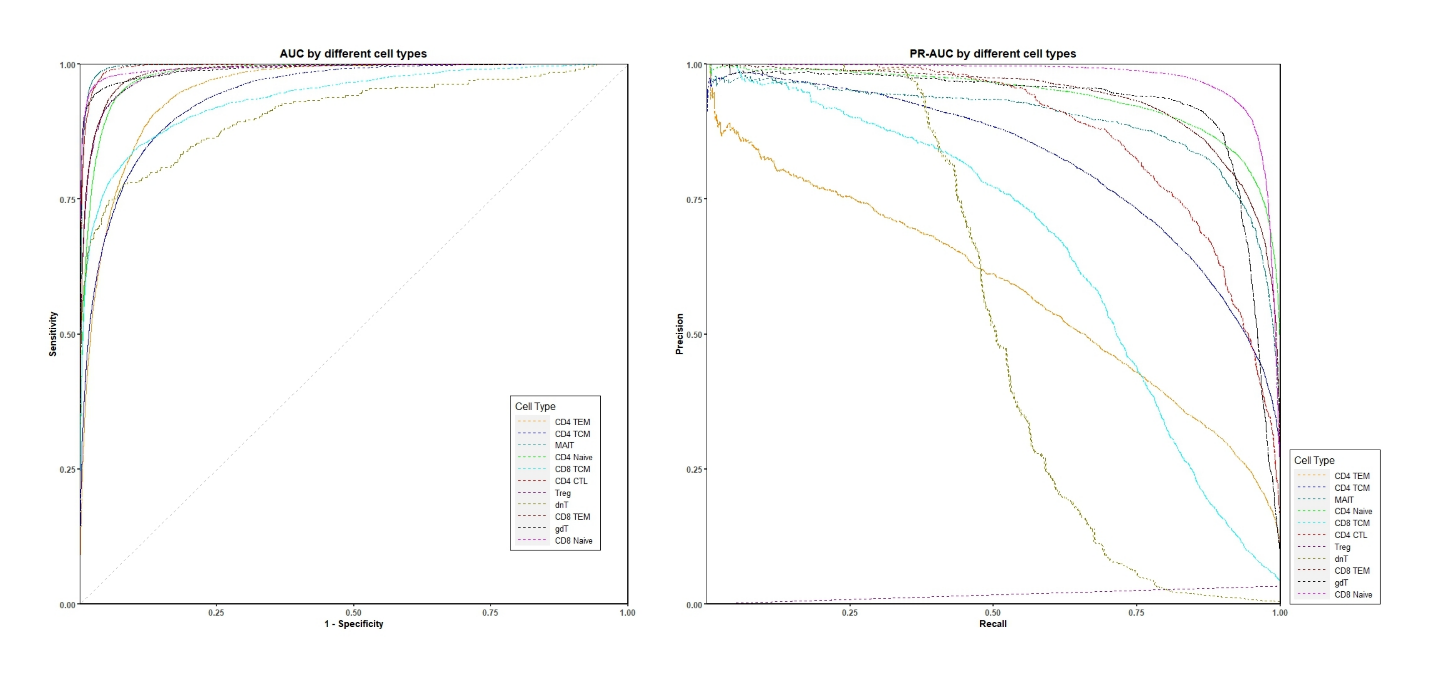

**Figure S6**. ROC (left) and PR-ROC (right) plots by different cell types on PBMC datasets (XGBoost)
